# Malaria parasites utilize pyrophosphate to fuel an essential proton pump in the ring stage and the transition to trophozoite stage

**DOI:** 10.1101/2021.10.25.465524

**Authors:** Omobukola Solebo, Liqin Ling, Jing Zhou, Tian-Min Fu, Hangjun Ke

## Abstract

The malaria parasite relies on anaerobic glycolysis for energy supply when growing inside RBCs as its mitochondrion does not produce ATP. The ring stage lasts ∼ 22 hours and is traditionally thought to be metabolically quiescent. However, recent studies show that the ring stage is active for several energy-costly processes including gene transcription/translation, protein export, and movement inside the RBC. It has remained unclear if a low glycolytic flux can meet the energy demand of the ring stage. Here we show that the metabolic by-product, pyrophosphate, is a critical energy source for the development of the ring stage and its transition to the trophozoite stage. During early phases of the asexual development, the parasite utilizes *Plasmodium falciparum* vacuolar pyrophosphatase 1 (PfVP1), an ancient pyrophosphate-driven proton pump, to pump protons across the parasite plasma membrane to maintain the membrane potential and cytosolic pH. Conditional deletion of PfVP1 leads to delayed ring stage development and a complete blockage of the ring-to-trophozoite transition, which can be partially rescued by *Arabidopsis thaliana* vacuolar pyrophosphatase 1, but not by the soluble pyrophosphatase from *Saccharomyces cerevisiae*. Proton-pumping pyrophosphatases are absent in humans, which highlights the possibility of developing highly selective VP1 inhibitors against the malaria parasite.

## Introduction

Malaria is a threat to 40% of the world’s population and claims more than 620,000 lives in 2020^1^. In a human host, the malaria parasite grows exponentially in bloodstream RBCs, causing all clinical symptoms including death in severe cases. In *Plasmodium falciparum*, the 48 h Intraerythrocytic Development Cycle (IDC) can be divided into three major developmental stages, including the ring, the trophozoite, and the schizont. These stages take about ∼ 22h, ∼ 18h, and ∼ 8h, respectively. Within the RBC, the parasite resides in a vacuole and is surrounded by three membranes: parasite plasma membrane (PPM), the parasitophorous vacuolar membrane (PVM), and the RBC membrane (RBCM). A major task of the ring stage parasite is to export proteins to the host cell to increase its permeability and cytoadherence^2^. Over the ∼ 22 h period, however, the parasite is not replicating DNA or expanding its biomass significantly. After the RBCM has been permeabilized by the Plasmodium Surface Anion Channel (PSAC)^3^, or New Permeability Pathways (NPPs)^4^, the trophozoite stage parasite starts to grow rapidly, resulting in 16-32 progeny in the subsequent schizont stage.

It has been long recognized that the asexual stage parasites rely on anaerobic glycolysis for ATP production^5,6^. Per glucose consumed, the parasite makes 2 ATP and 2 lactate molecules, with a minimal number of glucose-derived carbons fed into the tricarboxylic acid cycle (TCA)^7^. Indeed, the parasite can tolerate deletions of many TCA cycle enzymes^7^ and some components of the mitochondrial electron transport chain^8,9^, implying that the mitochondrion is a negligible source of ATP in blood stages. To overcome the energy constrain mediated by substrate-level phosphorylation, the trophozoite stage parasite runs a high rate of glycolysis and consumes glucose in a rate that is 100-times faster than normal RBCs^10^. Permeabilization of the RBCM in this stage also facilitates lactate disposal to avoid a metabolic blockage of glycolysis. With an intact RBCM, however, the ring stage is traditionally thought to be metabolically quiescent, with a low-level of glycolysis being sufficient to meet the energy demand of this stage^11^. Recent studies, however, suggest that the ring stage parasite fulfills many energy-costly processes over the 22 h period. Although the genome is not replicating at this stage, RNA transcription and protein translation are active to form a ring stage specific proteome for all necessary activities^12^. The PTEX translocon catalyzes ATP hydrolysis to move hundreds of parasite proteins to the RBC cytosol and membrane throughout the ring stage^13^. Rather than being static, ring stage parasites undergo dynamic movement inside the RBC and display morphological changes between the classical ring and a deformable ameboid-like structure^14^. In addition, the ring stage parasite must spend energy to pump protons across the parasite plasma membrane to maintain the plasma membrane potential (Δψ). It has been shown that the ATP-consuming V-type ATPase is the major proton pump in trophozoite stage parasites^15^. However, no studies have been carried out to show how plasma membrane potential is maintained in ring stages. RNA-seq data suggest that subunits of V-type ATPase are not highly transcribed until the trophozoite stage^16^ (***Figure 1-figure supplement 1***). Thus, it remains unknown how the ring stage parasite pumps protons and meets its energy demand while running a low-level of glycolysis.

In this study, we discover that the ATP independent, proton pumping pyrophosphatase PfVP1 (*<u>P</u>lasmodium falciparum* <u>v</u>acuolar <u>p</u>yrophosphatase 1), is the major proton pump during ring stage development. Proton-pumping pyrophosphatases, or H^+^-PPases, catalyze the hydrolysis of inorganic pyrophosphate (PPi), a by-product of over 200 cellular reactions, while harnessing the energy to pump protons across a biological membrane^17^. H^+^-PPase was first discovered in the plant tonoplast and was also named vacuolar pyrophosphatase^18^. While H^+^-PPases are absent in fungi and metazoans, it has been evolutionally conserved in bacteria, archaea, plants, and many protozoans^19^. The *P. falciparum* genome encodes two types of H^+^-PPases, PfVP1 (PF3D7_1456800) and PfVP2 (PF3D7_1235200)^20^. PfVP1 is potassium dependent and calcium independent whereas PfVP2 is potassium independent and calcium dependent. RNA-seq data suggest PfVP2 is barely transcribed^16^, which is consistent with its non-essential role in the asexual stages^21^; by contrast, PfVP1 is highly expressed throughout the IDC and exhibits a peak expression in the ring stage^16^ (***Figure 1-figure supplement 1***). Our data reveal that the malaria parasite employs PfVP1 to harness energy from pyrophosphate, an ancient energy source, to support vital biological processes in the ring stage when ATP supply is likely low.

## Results

### PfVP1 is mainly localized to the parasite plasma membrane (PPM)

Previous attempts of localizing vacuolar pyrophosphatases in *P. falciparum* used polyclonal antibodies raised against *Arabidopsis thaliana* vacuolar pyrophosphatase 1 (AVP1) in wildtype parasites^22^, which were unable to differentiate PfVP1 from PfVP2. Therefore, to specifically localize PfVP1, we utilized the CRISPR/Cas9 system^23,24^ to endogenously tag *pfvp1* with either a triple hemagglutinin (3HA) tag or a monomeric fluorescent protein (mNeonGreen) in the 3D7-PfVP2KO (knockout) parasite line^21^. Additionally, through gene editing of the endogenous copy, the tagged *pfvp1* was placed under the control of the TetR-DOZI-aptamer system for conditional expression^25,26^ (***Figure 1- figure supplement 2***). Thus, two transgenic parasite lines were constructed, 3D7-PfVP2KO-PfVP1-3HA^apt^ and 3D7-PfVP2KO-PfVP1-mNeonGreen^apt^. We also cloned 3D7-PfVP2KO-PfVP1-3HA^apt^ by limited dilution and obtained two pure parasite clones, B11 and G11, which were phenotypically indistinguishable (B11 was used for this study). The parasite lines were normally cultured in the presence of 250 nM anhydrotetracycline (aTc) to maintain PfVP1 expression.

The subcellular localization of PfVP1 was verified by immunofluorescence analysis (IFA), immuno-electron microscopy (immuno-EM), and live fluorescence microscopy (***Figure 1***). In the 3D7-PfVP2KO-PfVP1-3HA^apt^ line, IFA revealed clear colocalization of PfVP1 and the PVM marker, PfEXP2, throughout the 48 h IDC (***Figure 1A***). Since the PVM is permeable to protons^27^, the close proximity of PfVP1 to PfEXP2 suggests PfVP1 is localized on the PPM. Further, immuno-EM studies with the HA tagged line showed localization of PfVP1 mainly to the PPM (***Figure 1B***). Quantification of 65 random images revealed 90% of the gold particles were localized to the PPM, ∼ 2% of the gold particles were localized to nucleus/ER, and ∼ 8% of the gold particles were localized to the cytosol or cytosolic small membranous structures with unknown identities (***Figure 1- figure supplement* 3**). No gold particles were apparently localized on the food vacuole. To further confirm this, we performed co-localization studies of PfVP1 and the food vacuole using the food vacuole marker, PfPlasmepsin II^28^. We were unable to find any parasites in which PfVP1 and PfPlasmepsin II colocalized (***Figure 1- figure supplement* 4**). Thus, PfVP1 did not coincide with the food vacuole, as previously thought^29^. Finally, we used live microscopy to localize PfVP1 in the 3D7-PfVP2KO-PfVP1-mNeonGreen^apt^ line. PfVP1 was clearly localized to the PPM in every stage of the IDC, including the merozoite, the ring, the trophozoite, and the schizont stages (***Figure 1C and D***). Together, we utilized three independent methods to show that PfVP1 is mainly localized to the PPM.

**Figure 1.**
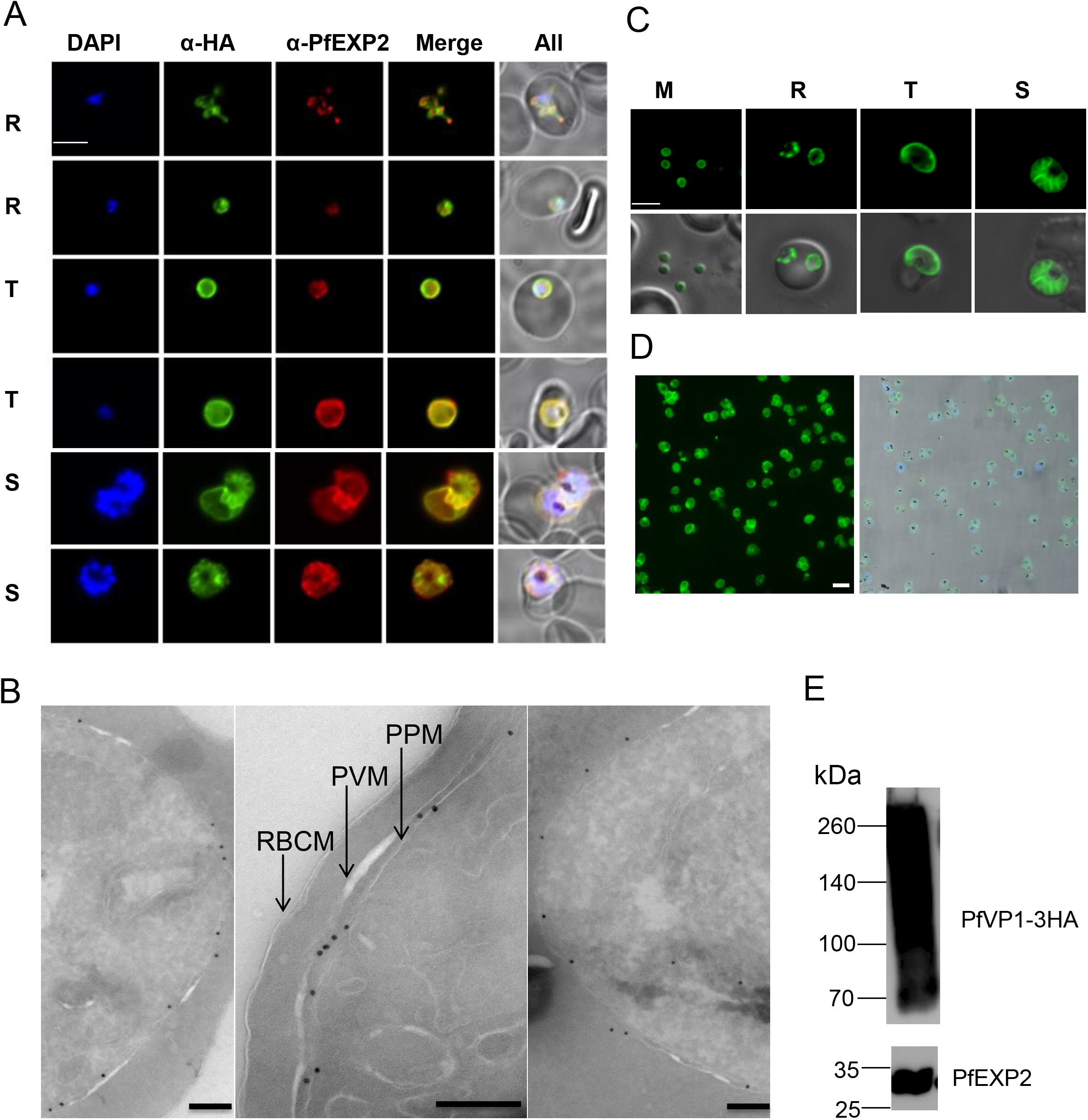
PfVP1 is mainly localized to the parasite plasma membrane. A, Immunofluorescence assay of Pf3D7VP2KO-VP1-3HA^apt^. DAPI stains the nuclei. Green, PfVP1-3HA. Red, PfEXP2. Note an ameboid ring stage parasite in the first row. R, ring. T, trophozoite. S, schizont. Scale bar, 5 µm. Representative images of n>50 parasites of each stage are shown here. B, Immunoelectron microscopy of Pf3D7VP2KO-VP1-3HA^apt^. RBCM, RBC membrane. PVM, parasitophorous vacuolar membrane. PPM, parasite plasma membrane. Scale bars, 200 nm. C, Live imaging of Pf3D7VP2KO-VP1-mNeonGreen^apt^. M, merozoite. R, ring. T, trophozoite. S, schizont. Scale bar, 5 µm. D, Live imaging of Percoll enriched Pf3D7VP2KO-VP1-mNeonGreen parasites. Scale bar, 10 µm. E, Western blot showing PfVP1-3HA expression. The blot was re-probed with anti-PfExp2 antibody to show the loading control. Approximately, 15 µg of total protein lysate was loaded in the gel. The online version of this article has the following figure supplements for figure 1. Figure supplement 1. RNA-seq data of pyrophosphatases and proton pumps in *P. falciparum*. Figure supplement 2. Genetic tagging of PfVP1 via CRISPR/Cas9. Figure supplement 3. Localization of PfVP1 via Immunoelectron microscopy. Figure supplement 4. Co-localization of PfVP1 and Plasmepsin II via Immunofluorescence assay.

In the 3D7-PfVP2KO-PfVP1-3HA^apt^ line, we revealed PfVP1 was also highly expressed by Western blot (***Figure 1E***). A regular amount of total parasite lysate (∼ 15 µg) contained an abundant amount of PfVP1, from the monomeric form of ∼ 79 kDa to large, aggregated oligomers that were not apparently solubilized by 2% SDS. The aggregated forms agreed with the fact that PfVP1 contains 16 transmembrane helices and is highly insoluble.

### Characterizing PfVP1 using the Saccharomyces cerevisiae heterologous system

To confirm PfVP1 is a PPi-dependent proton pump, we expressed PfVP1 in *S. cerevisiae*. Since the 1990s, this heterologous system^30^ has been widely applied to study many VP1 orthologs from plants and Archaea^31-33^. *S. cerevisiae* does not have VP1 homologs and thus provides a robust and clean system to study exogenous VP1 proteins^30^. Importantly, isolated yeast vacuolar vesicles incorporating recombinant VP1 are suitable for testing the pump’s ability to move protons from one side of the membrane to the other. The vesicles can also be used to examine VP1’s enzymatic activity. We transformed the yeast strain BJ5459^34,35^, which was null for the two major vacuolar proteases, PrA and PrB, with plasmids containing a copy of synthetic codon optimized PfVP1, AVP1, or a blank control. VP1 proteins were N-terminally tagged with the localization peptide of *Trypanosoma cruzi* VP1 (the first 28 amino acids) and GFP, which facilitates VP1’s localization to yeast vacuoles^36^. Yeast expression of PfVP1 and AVP1 was verified by fluorescence microscopy, which showed that the GFP signal appeared mainly on the yeast vacuoles (***Figure 2-figure supplement 5***).

We followed the established protocols^37^ to isolate yeast vesicles from these three lines expressing PfVP1, AVP1, or the blank control (***Figure 2A***). In isolated yeast vesicles, a 9-Amino-6-Chloro-2-Methoxyacridine (ACMA) fluorescence quenching assay was used to assess the ability of VP1 to pump protons into their lumen (Materials and Methods). The compound’s fluorescence is quenched when a pH gradient forms across the vesicle membrane. The yeast V-type ATPase, also present on the vesicles, was inhibited by Bafilomycin A1. Over time, PfVP1 possessing vesicles were able to reduce ACMA fluorescence (***Figure 2B***). The positive control AVP1 expressing vesicles also quenched ACMA, as expected, whereas the negative control vesicles bearing no H^+^-PPase exhibited little effect. When Nigericin was added (a proton ionophore that abolishes transmembrane proton gradients), the quenched ACMA fluorescence was restored to its original levels (***Figure 2B***). This verified that the yeast vesicles were intact and PfVP1 and AVP1 expressing vesicles had accumulated protons inside. We also assessed PfVP1’s enzymatic activity by measuring free Pi released by PPi hydrolysis (Materials and Methods). In comparison to the negative control vesicles, PfVP1 expressing vesicles produced a net of 1.24 µmoles free Pi per mg of protein per h, similar to that produced by the AVP1 vesicles (1.14 µmoles/mg/h) (***Figure 2C***). Together, using the yeast heterologous expression system, we confirmed that PfVP1 is a PPi hydrolyzing proton pump.

**Figure 2.**
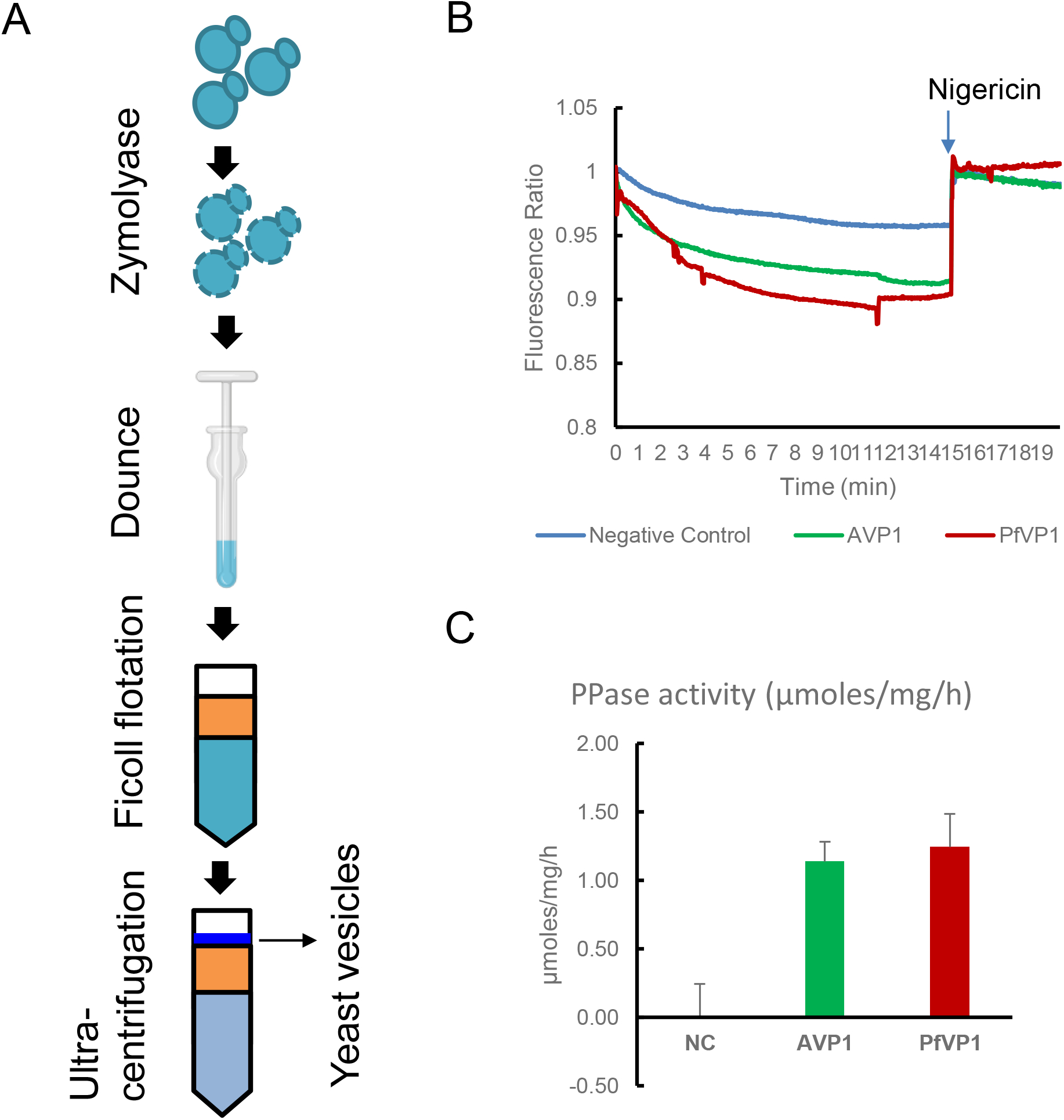
PfVP1 is a PPi hydrolyzing proton pump. A, A general schematic of purifying yeast vesicles bearing VP1 proteins from *Saccharomyces cerevisiae*. Yeast cells were treated with Zymolyase to remove the cell wall, lysed by Dounce, and applied to a ficoll gradient (16% and 8%). After ultracentrifugation, yeast vesicles were collected from the top. B, ACMA quenching assay in isolated yeast vesicles bearing AVP1(*A. thaliana* vacuolar pyrophosphatase 1), PfVP1, or no VP1 protein (negative control). The ACMA’s fluorescence signal was recorded after the yeast vesicles were added with the substrate, Na_2_PPi. Data shown are the representative of five individual experiments. C, Pyrophosphatase activity measurement in isolated yeast vesicles bearing AVP1, PfVP1, or no VP1 protein. The background activity from the negative control (NC) vesicles was subtracted from all measurements. This experiment was repeated three times with technical replicates. Data shown here are the mean ± standard deviations of the replicates. The online version of this article has the following figure supplement for figure 2. Figure supplement 5. Localization of VP1 proteins in *S. cerevisiae*.

### PfVP1 is essential for ring stage development and its transition to trophozoite

To investigate PfVP1’s essentiality during the 48h IDC, we set up knockdown studies and examined parasite viability and morphology in the 3D7-PfVP2KO-PfVP1-3HA^apt^ line. We used two approaches to remove aTc from cultures to initiate PfVP1 knockdown. In one approach, aTc was removed from Percoll isolated schizonts. In the other, aTc was removed from synchronized ring stage parasites. When aTc was removed from schizonts, the knockdown culture did not display discernible changes in parasite morphology or parasitemia in the first IDC (***Figure 3A***). This was likely because a prolonged time (∼ 48 h) was needed to knock down greater than 95% of the PfVP1 protein (***Figure 3-figure supplement 6A***). However, in the second IDC after aTc removal, the knockdown parasites then showed drastic morphological changes (***Figure 3A***). Although invasion was apparently unaffected in the absence of PfVP1 (***Figure 3-figure supplement 6B***), the parasites struggled to progress through the ring stage and failed to become an early stage trophozoite in the second IDC. From 72 to 96 h, while the parasites in aTc (+) medium progressed normally from the ring to schizont stage, the size of the knockdown parasites barely increased. It appeared that PfVP1 knockdown resulted in an extended ring stage as long as ∼ 48 h. To better understand the morphological changes in the 2^nd^ IDC after aTc removal, we examined parasite development every 4 h (***Figure 3B***). Again, PfVP1 knockdown caused delayed ring stage development and a complete blockage of the ring to trophozoite transition. At the end of the 2^nd^ IDC, we noticed that the knockdown parasites had expanded the cytosol slightly in comparison to parasites of earlier time-points and small hemozoin particles were also visible (***Figure 3B***). However, none of the knockdown parasites showed the morphology of a normal early trophozoite stage parasite. In the absence of PfVP1, the parasites were arrested in this state for several days before lysing and was unable to finish the asexual cycle. Moreover, the blockage of ring to trophozoite transition was also observed when the knockdown experiment was initiated from ring stage parasites (***Figure 3C***).

**Figure 3.**
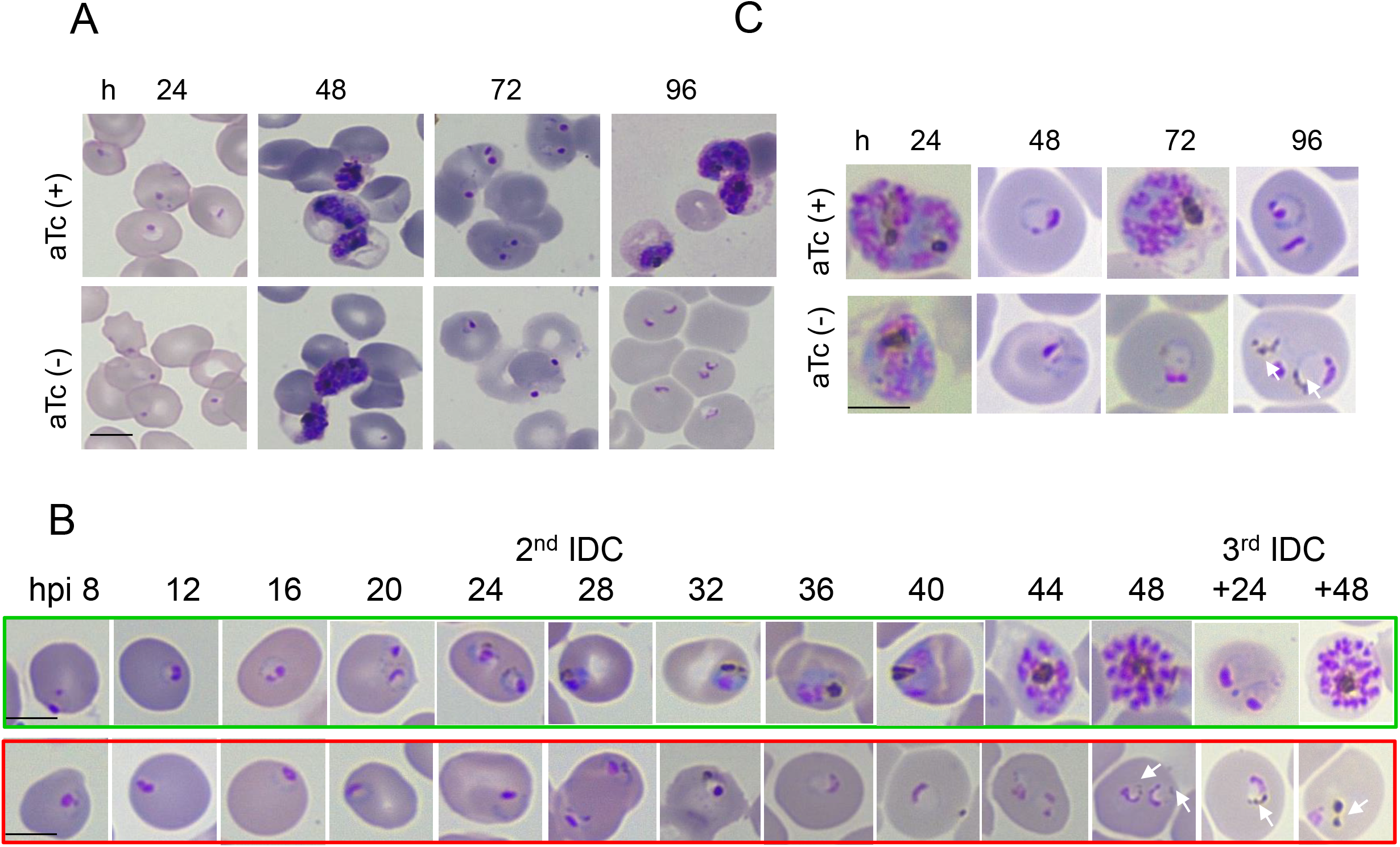
PfVP1 is essential for the ring stage and its transition to trophozoite stage. A, Knockdown experiment starting at the schizont stage by removal of aTc. B, Parasite morphological changes throughout the time course. Green box, aTc (+). Red box, aTc (-). C. Knockdown experiment starting at the ring stage by removal of aTc. A-C, images were Giemsa-stained thin blood smears taken by a light microscope. Bars, 5 µm. B-C, White arrows indicate small hemozoin particles. These experiments were repeated more than five times (A, C) or two times (B). The online version of this article has the following figure supplements for figure 3. Figure supplement 6. PfVP1 is highly expressed and essential for the ring to trophozoite transition. Figure supplement 7. Arrested ring stage parasites can poorly progress when aTc is restored.

To examine if the arrested parasites are viable, we added aTc back to the culture after aTc was removed for 72, 84 and 96 h from schizonts. The arrested ring stage parasites were able to fully progress if aTc was given back at 72 and 84 h post knockdown (data not shown), indicating that they remained viable. When aTc was added back at 96 h after knockdown, however, the arrested parasites displayed poor growth in the next IDC. Although most of them were able to progress to the trophozoite stage, many appeared sick-looking and lysed on the next day (***Figure 3-figure supplement 7A***). The total parasitemia only increased marginally in the aTc addback culture (***Figure 3-figure supplement 7B***). Altogether, this data clearly establish that PfVP1 is essential for ring stage development, and its absence halts the maturation of the parasites at the late ring stage.

### Phenotypic characterization of the PfVP1 knockdown parasites

We next characterized the knockdown phenotypes in the 3D7-PfVP2KO-PfVP1-3HA^apt^ line. We reasoned if PfVP1 was genuinely a PPi-dependent proton pump located on the PPM, changes to cytosolic pH and PPi levels would be expected in the knockdown parasite. We used the ratiometric pH sensitive dye 2’,7’-Bis-(2-Carboxyethyl)-5-(and-6)-Carboxyfluorescein, Acetoxymethyl Ester (BCECF-AM), to measure cytosolic pH following the established protocol^19^. BCECF-AM is membrane permeable and is trapped inside the parasite cytosol after its ester group is removed. We initially aimed to measure cytosolic pH at 48, 72 and 96 h after aTc removal from schizonts; however, attempts to measure pH in ring stage parasites were unsuccessful. Our experience agreed with the fact that BCECF-AM has only been reported to measure cytosolic pH in trophozoite stage parasites^38,39^, not in the ring stage. Nonetheless, we successfully detected a decrease of cytosolic pH in the knockdown culture at 48 h post aTc removal (***Figure 4A***), although the knockdown parasites remained morphologically healthy. Next, we measured PPi levels in the knockdown cultures using a newly developed PPi specific sensor (Materials and Methods). At 72, 84, and 96 h post aTc removal, saponin-lysed pellets were collected, and soluble metabolites were extracted using a mild process (Materials and Methods). In each sample, the concentrations of PPi and total parasite protein were measured, and the total amount of PPi (nanomoles) was normalized to total protein (mg). We observed an increase of PPi at 84 and 96 h after aTc removal (***Figure 4B***). Together, these data not only reveal the mode of action of PfVP1 in *Plasmodium*, but also suggest that when PfVP1 is knocked down, the cytosolic proton and PPi levels increase, likely preventing parasite’s transition from the ring to trophozoite stage.

**Figure 4.**
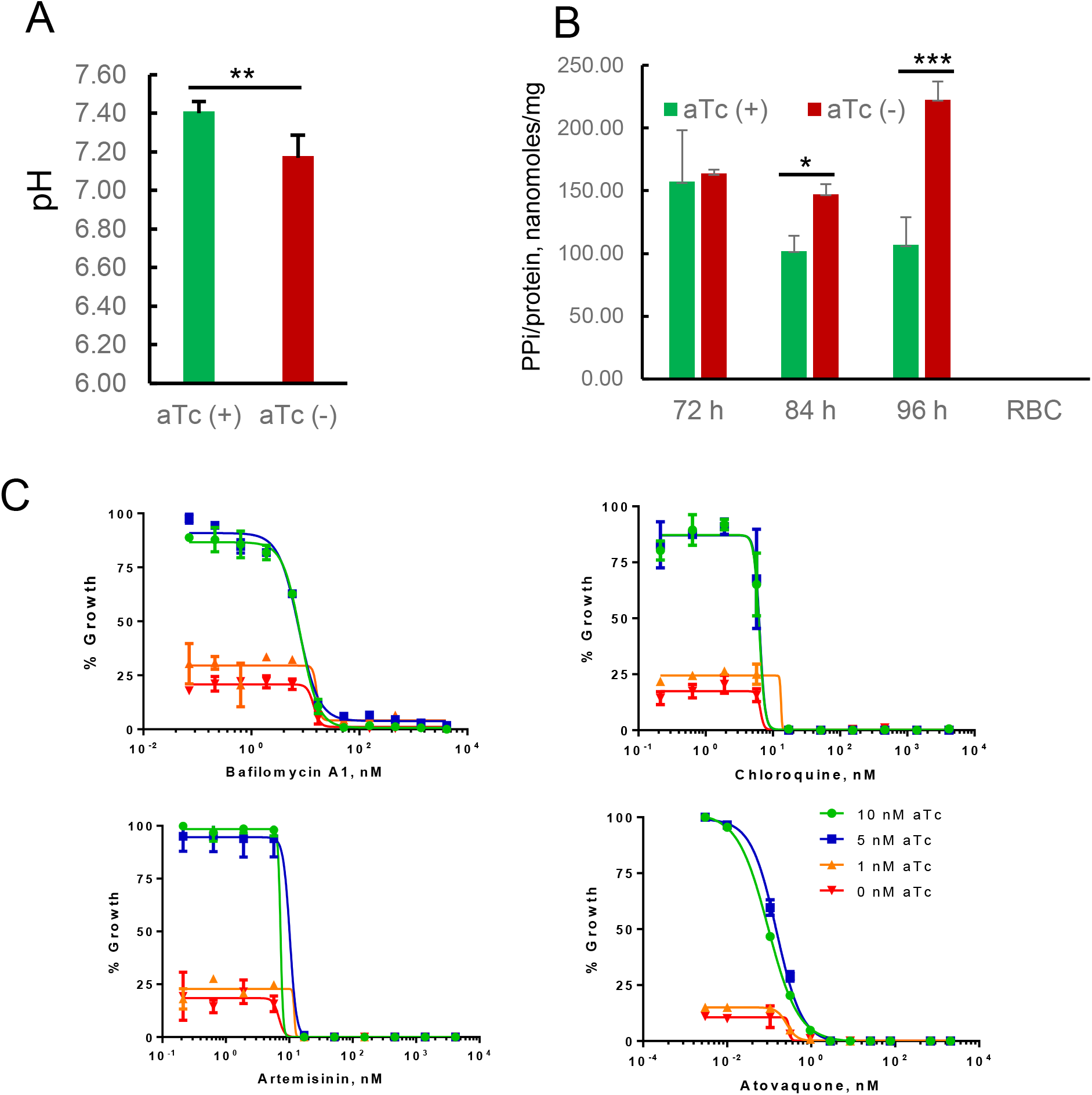
Phenotypic analysis of the PfVP1 knockdown parasite. A, pH measurement in the knockdown parasites after aTc removal for 48 h from the schizont stage. B, PPi measurement in the knockdown parasites after aTc removal for 72, 84, and 96 h from the schizont stage. In A-B, error bars indicate standard deviation of n=3 measurements in each condition; statistical analysis was done by Student t-test. *, *p* < 0.05. **, *p* < 0.01. ***, *p* < 0.001. Experiments of A and B were repeated twice. C, Sensitivity to antimalarials measured by SYBR green assays. EC_50_ values were only retrievable from cultures grown in 10 or 5 nM aTc, as listed below: Bafilomycin A1 (8.1 vs 7.5 nM), Chloroquine (6.3 vs 6.4 nM), Artemisinin (7.3 vs 10.2 nM), Atovaquone (0.1 vs 0.15 nM). This experiment was repeated three times.

We next employed a chemical-genetic approach to examine how the knockdown parasites responded to antimalarials. Starting at ring stages, we washed out aTc and set up 72h SYBR green assays with varying concentrations of aTc (10, 5, 1 and 0 nM). We used 10 nM as the highest concentration since it was sufficient to support 100% parasite growth (data not shown). Drug inhibition plots and EC_50_ values obtained with parasites grown at 10 or 5 nM aTc were similar to those found with wild type parasites. However, the PfVP1 knockdown parasites became hypersensitive to all antimalarials tested when aTc concentrations were reduced to 1 or 0 nM (***Figure 4C***). This data indicated that when PfVP1 was knocked down, the parasite was so ill that it became sensitive to extremely low concentrations of all antimalarials tested. Inhibitors against PfVP1, if available, would therefore be expected to have synergy with many antimalarials. Of note, efforts to develop PfVP1 inhibitors have already begun by others^40^.

### Complementation of the knockdown parasite line with yIPP1 or AVP1

In *A. thaliana*, AVP1 acidifies the plant vacuole in conjunction with V-type ATPase. The proton pumping activity of AVP1 is not as critical as the PPi hydrolysis activity since a loss-of-function of AVP1 was rescued by the yeast inorganic pyrophosphatase (yIPP1), the soluble pyrophosphatase from *S. cerevisiae*^41^. yIPP1 has the sole function of PPi removal, with no energy saving or proton pumping activity. Therefore, the proton pumping activity and PPi hydrolysis activity of AVP1 can be de-coupled. To test if this is also true for PfVP1, we performed a second round of transfection to complement the knockdown parasite with a copy of Myc-tagged yIPP1, AVP1 or wildtype PfVP1.

To this end, we made a new knockdown line in the D10 wildtype background, resulting D10-PfVP1-3HA^apt^ line. We transfected D10-PfVP1-3HA^apt^ parasites with plasmids bearing hDHFR for selection^42^ and 3Myc-tagged yIPP1, AVP1 or PfVP1(Materials and Methods). Western blots showed that all Myc tagged copies were expressed independent of aTc, while the endogenous HA tagged PfVP1 was knocked down when aTc was removed for 96 h (***Figure 5A and B***). As expected, fluorescence microscopy showed PfVP1 or AVP1 was localized to the PPM whereas yIPP1 was in the cytosol (***Figure 5C***). When aTc was removed from schizonts for 96 h, the knockdown parasite complemented with PfVP1-3Myc displayed normal growth like the aTc (+) control, indicating that the episomal PfVP1 fully complemented the endogenous copy (***Figure 5D***). AVP1 complementation displayed a moderate rescue with two thirds of the parasites reaching the same morphology as control parasites (late trophozoite), and one third progressing to a smaller size (early trophozoite). In contrast, yIPP1 was unable to restore parasite growth when the endogenous PfVP1 was knocked down. A quantification of various parasite morphologies in all conditions is shown in ***Figure 5E***. Altogether, this data indicates 1) both PfVP1’s PPi hydrolysis and proton pumping activities are essential for parasite survival, and 2) although not a 100% functional replacement, the homologous plant VP1 is able to complement VP1-deficient *Plasmodium* parasites.

**Figure 5.**
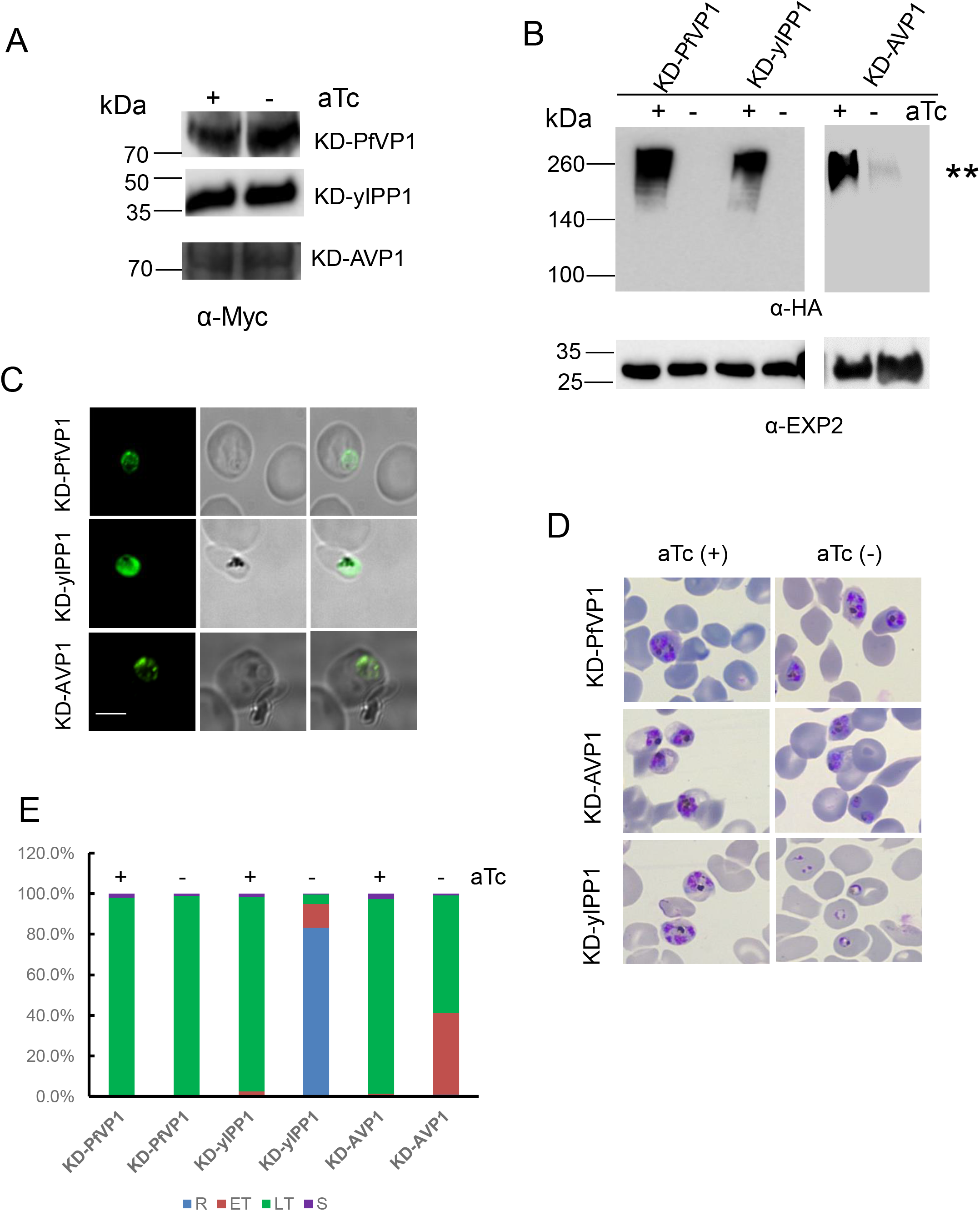
The dual functionality of PfVP1 is required for parasite survival. A, Western blot analysis of protein lysates from the D10-PfVP1-3HA^apt^ line that was episomally transfected with Myc tagged PfVP1 (wildtype), yIPP1 (yeast inorganic pyrophosphatase), or AVP1 (*A. thaliana* vacuolar pyrophosphatase 1). Parasites were grown in aTc (+) or (-) conditions for 2 days. Blots were probed with anti-Myc antibody and its secondary antibody. B, Western blot analysis of protein lysates from the D10-PfVP1-3HA^apt^ line that was episomally transfected with Myc tagged PfVP1, yIPP1, or AVP1. Parasites were grown in aTc (+) or (-) conditions for 2 days. Approximately, ∼ 3 µg of total protein lysate from each line was loaded in the gels. Blots were first probed with anti-HA antibody to show the expression levels of the endogenously tagged PfVP1 with 3HA. **, when a small amount of total protein lysate was loaded, only aggregated forms of PfVP1 with high molecular weights were detected by Western blot. The same blots were re-probed with anti-PfExp2 antibody to show loading controls. C, Immunofluorescence analysis (IFA). The complemented lines were probed with anti-Myc and a fluorescent secondary antibody. Scale bar, 5 µm. Representative images of n>30 parasites of each line were shown here. D, Morphologies of complemented parasite lines at 96 h after aTc removal. This experiment was repeated three times. E, Quantification of parasite morphologies in D. The percentage of different parasite morphological stages was determined by counting 1000 infected RBCs in each condition. R, ring. ET, early trophozoite. LT, late trophozoite. S, schizont. This experiment was repeated two times. A-E, KD means knockdown.

### Structure-guided mutagenesis studies of PfVP1

To further understand the mode of action of PfVP1, we conducted structure-guided mutagenesis studies in *P. falciparum*. All VP1 orthologs have 15-17 transmembrane helices with a molecular mass of 70-81 kDa^19^. The crystal structure of *Vigna radiata* (mung bean) VP1 (VrVP1) was resolved in 2012^17^. At the primary sequence level, PfVP1 is highly similar to VrVP1 (49% identity and 66% similarity). The transmembrane (TM) helices are well conserved between PfVP1 and VrVP1, although the inter-domain loops display noticeable differences (***Figure 6A***). VrVP1 contains longer loops between the first three TMs. Based on the crystal structure, we computationally modeled the structure of PfVP1; the model showed a high degree of conservation to VrVP1 with deviations in some loop regions (***Figure 6B***). The substrate binding and hydrolyzing site of the modeled PfVP1 also mimics that of VrVP1^17^. At this site, all the conserved residues including 8 aspartates and 1 lysine are positioned around the substrate analog, the magnesium imidodiphosphate (MgIDP) (***Figure 6C***). The proton transfer pathway formed by TMs 5, 6, 11, 12 and 16 also appears to be structurally conserved (***Figure 6D***).

**Figure 6.**
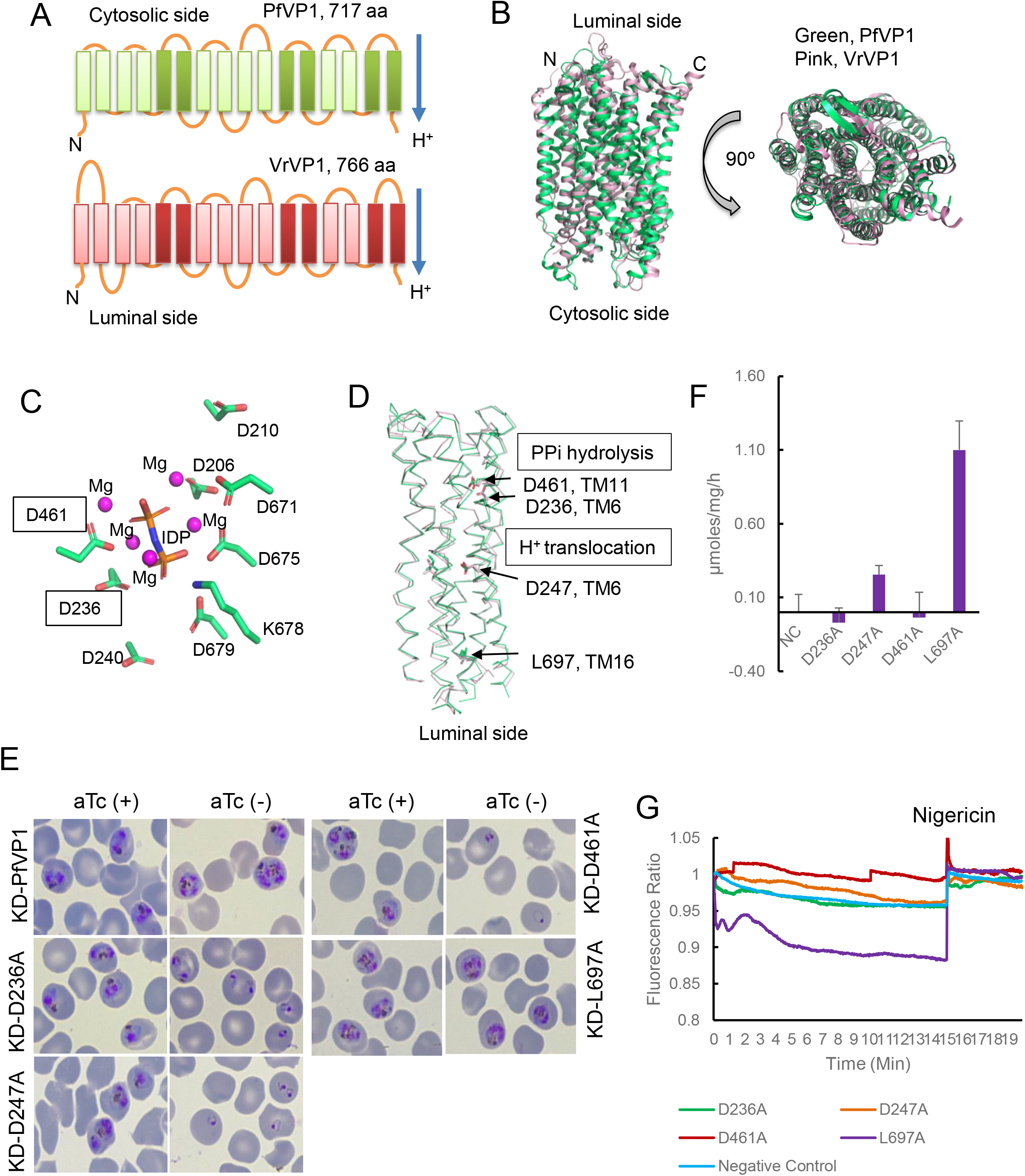
Structure guided mutagenesis analysis of PfVP1. A, 2D schematic of PfVP1 and *Vigna radiate* VP1 (VrVP1) containing 16 transmembrane helices (TMs). In each monomer, TMs of 5, 6, 11, 12, 15, 16 (darker color) form the inner circle whereas the rest 10 TMs (lighter color) form the outer circle. Protons are pumped from the cytosolic side to the luminal side. B. Structure of PfVP1 (green) overlayed with the crystal structure of VrVP1 (pink). The PfVP1 structure was predicted using RoseTTaFold. C. Substrate binding site of PfVP1. The side chains of substrate binding amino acids were highlighted in sticks. Magnesium ions were shown in magenta spheres. IDP stands for imidodiphosphate, which was used to co-crystalize VrVP1^17^. Boxed residues will be mutated. D. Side view of the inner circle formed by TM5, TM6, TM11, TM12 and TM16. The proton transfer pathway is located at the lower part of the inner circle. Residues subjected to mutagenesis are indicated. E, Parasite morphologies of mutated PfVP1 lines at 96 h after aTc removal from the schizont stage. Giemsa-stained smears were shown. This experiment was repeated three times. F, Pyrophosphatase activity measurement in isolated yeast vesicles bearing different mutant PfVP1 proteins. The background activity of negative control (NC) yeast vesicles was subtracted from all measurements. This experiment was repeated three times (n=3). Data shown here are the mean ± standard deviations of the replicates. G, Proton pumping activity of isolated yeast vesicles bearing different mutant PfVP1 proteins. Data shown are the representative of 3-5 experiments in each yeast line. The online version of this article has the following figure supplement for figure 6. Figure supplement 8. Localization and expression of PfVP1 mutant alleles.

Based on these structural analyses, we chose to do alanine replacement mutagenesis of two putative substrate binding residues (D236, D461) and two residues that appear to be in the proton transfer channel and exit gate (D247, L697). Since *Plasmodium* is haploid and direct mutagenesis of essential residues would be lethal, we performed these mutagenesis studies in the D10-PfVP1-3HA^apt^ line by episomal expression of mutated alleles (Materials and Methods). The effect of mutant PfVP1 on parasite viability was assessed upon knockdown of the endogenous copy by aTc removal. Fluorescence microscopy showed all mutant PfVP1 proteins were expressed and localized to the PPM (***Figure 6-figure supplement 8A***). When the endogenous HA tagged PfVP1 was knocked down by aTc removal for 96 h, all Myc tagged PfVP1 mutant alleles were still expressed (***Figure 6-figure supplement 8B***).

The mutant PfVP1 alleles had differing abilities to rescue the knockdown phenotype (***Figure 6E***). As a control, the episomal wildtype PfVP1 copy fully rescued the knockdown parasites grown in aTc (-) medium. PfVP1 alleles with D236A and D247A mutations were unable to rescue, indicating these mutations abolished the pump’s functions. The PfVP1/D461A mutation had partial rescuing ability, but most D461A expressing parasites were unable to progress to the trophozoite stage. These results largely agreed with the results obtained with equivalent mutations in VrVP1^43^. In contrast, PfVP1 differed from VrVP1 at mutation of L697A. When this residue in VrVP1 was mutated (L749A), the pump has lost proton pumping activity, although the PPi hydrolysis activity was largely remained^43^. However, the PfVP1/L697A allele had both PPi hydrolysis and proton pumping activities to fully rescue the knockdown culture. Since L697 is located at the proton exit gate, our results indicate that PfVP1 may have some structural variation from VrVP1 at least in this location. A quantification of the rescuing ability of the various mutant PfVP1 alleles is shown in ***Figure 6-figure supplement 8C***.

To explore the enzymatic and proton pumping activities of each mutant PfVP1 protein, we utilized the yeast heterologous expression system as described in ***Figure 2***. We individually purified yeast vesicles bearing different codon optimized PfVP1 mutant alleles from the BJ5459 strain. Fluorescence microscopy showed all modified PfVP1 proteins were expressed and localized to the yeast vacuole (***Figure 6-figure supplement 8D***). The enzymatic assays revealed that all mutant PfVP1s, except for the L697A mutation, lacked PPi hydrolysis activity (***Figure 6F***). Likewise, only PfVP1/L697A showed proton pumping activity in the ACMA quenching assay (***Figure 6G***). Altogether, using homology modeling, yeast and parasite expression systems, we find that the substrate binding site and proton transfer pathway of PfVP1 appear to be well conserved, although the proton exit gate of PfVP1 may differ from that of the plant VP1.

## Discussion

Our study has revealed that PfVP1 is highly expressed and mainly localized to the parasite plasma membrane (***Figure 1***). Using the yeast heterologous expression system, we demonstrated that PfVP1 is a PPi dependent proton pump (***Figure 2***). Our genetic data uncovered the essential role of PfVP1 in early phases of the IDC, including the ring stage and the ring to trophozoite transition (***Figure 3***). Conditional deletion of PfVP1 also resulted in the accumulation of protons and PPi in the parasite (***Figure 4***). Overall, our data indicate that the malaria parasite utilizes the ATP-independent proton pump PfVP1 to harness energy from pyrophosphate, a metabolic by-product, to establish the parasite plasma membrane potential (Δψ) in the ring stage.

While the IDCs of different malaria parasites vary between 24-72 h, the ring stage is invariably the longest period. In *P. falciparum*, the duration of the ring stage (∼ 22 h) combined with the transition stage from the ring to trophozoite (∼2-4 h) is half of the entire IDC^14^. Inside the RBC, the ring stage parasite moves, changes its shape^14^, and is busy exporting hundreds of proteins to the host cell^2^. Moreover, the ring stage is less susceptible to many antimalarial drugs and is the only stage that displays artemisinin resistance^44^. During the transition stage from the ring to trophozoite, the parasite also exhibits pronounced changes, including a reduction in the parasite diameter, formation of several small hemozoin foci, and a transient echinocytosis of the host cell (RBC membrane distortion)^14^. Despite the significance of these early phases of parasite development, little is known about their cellular bioenergetics.

Earlier studies have shown that the ring stage parasite performs glycolysis at a much lower rate compared to that of the trophozoite stage^11^. Traditionally, a low-level of glycolysis is thought to be sufficient to support ring stage development. However, our study has revealed that the metabolic by-product PPi serves as a critical energy source during the early phases of the IDC. The free energy of PPi hydrolysis under physiological conditions is estimated to be -22.18 kJ/mol, which is a significant portion of the energy released from ATP hydrolysis (−37.6 kJ/mol)^45^. Evolutionarily, early life forms on earth used PPi as the energy source before ATP emerged^45^. The early divergent malaria parasite has evolutionarily reserved the ability to use PPi as a critical energy source, especially at the time when the ATP level is low. During the IDC, the malaria parasite’s energy supply depends on inefficient ATP production via anaerobic glycolysis as the parasite’s mitochondrion is not performing oxidative phosphorylation. Therefore, PPi becomes a significant energy supplement to the ring stage parasite where glycolysis runs at a lower rate. Future studies will focus on quantifications of ATP and PPi throughout the IDC to understand their energetic contributions to the malaria parasite.

Unlike many other eukaryotes, malaria parasites generate the plasma membrane potential (Δψ) through the transport of protons rather than sodium ions^46^. The proton gradient across the plasma membrane is also used by the parasite to perform secondary active transport to move ions, nutrients, or waste products into or out of the cell^47^. It has been long recognized that the malaria parasite possesses two different types of proton pumps, the single subunit PPi-dependent H^+^-PPases^20,22^ and the much faster ATP-dependent multi-subunit V-type ATPase^48^. Inhibition of the V-type ATPase by Bafilomycin A1 for 10-12 min causes a rapid drop of cytosolic pH from ∼ 7.3 to ∼ 6.8 in trophozoite stage parasites^15^. Therefore, the much slower proton pumps, PfVP1 and PfVP2, were thought to be insignificant or “marginal” to the parasite^48^. Alternatively, other studies have hypothesized that PfVP1 and/or PfVP2 would be critical to the parasite when energy demand is high in trophozoite stage parasites^20^. In contrast to those earlier views, our results have now recognized the significance of PfVP1 for ring stage development (PfVP2 is dispensable for asexual development^21^). Our data suggest that by using PfVP1 to fulfill proton pumping across the PPM, the ring stage parasite can divert ATP to other energy costly processes such as protein export. Therefore, for the first time, we have shown that PfVP1 is the major proton pump in *Plasmodium falciparum* during the ring stage development. Without PfVP1, the parasite is arrested at the late ring stage and fails to become an early trophozoite.

It is interesting to note that the ortholog VP1 protein in *Toxoplasma gondii* (TgVP1) displays different subcellular localization and function. TgVP1 is mainly localized to acidocalcisomes and the plant-like vacuole (PLV)^49,50^ and despite phenotypic alterations, a complete knockout of TgVP1 is tolerated by the parasite^50^. In *P. falciparum*, however, the large-scale mutagenesis survey was unable to disrupt the PfVP1 gene^51^. We have shown here that PfVP1 is essential for the ring stage development. Hence, the conserved VP1 protein has seemingly adapted to perform different functions even within the Apicomplexa phylum, to which *Toxoplasma* and *Plasmodium* belong to. Unlike blood stage malaria parasites, *Toxoplasma gondii* has a much more robust mitochondrion that is a significant ATP producer^52^. On the other hand, our study has not ruled out the possible localization of PfVP1 to acidocalcisomes in malaria parasites. In contrast to *T. gondii* where acidocalcisomes are more abundant, the presence of these organelles in *Plasmodium* remains obscure. Except for merozoites^53^, the classical acidocalcisomes have not been firmly reported in the literature in other asexual stage parasites. Further investigation is needed to clarify if acidocalcisomes are present in malaria parasites and if PfVP1 is present on them.

In summary, our data suggest that the malaria parasite utilizes PfVP1 to harness energy from PPi to establish the plasma membrane potential, extrude cytosolic protons, and maintain an energy homeostasis in early development of the IDC. The essential nature of PfVP1 combined with the absence of any orthologs in humans has also highlighted it as a potential antimalarial drug target. A drug target in the ring stage is highly desired to the drug development pipeline, inhibitors of which could be partnered with many other antimalarials that kill metabolically more active stages. A combination therapy targeting both young and mature malaria parasites can then ensure all parasite forms are dispatched. Although at early stages, efforts of developing inhibitors against H^*+*^-PPases including PfVP1 have already begun^40^.

## Materials and Methods

### 1, Plasmid construction for *P. falciparum* studies

#### 1) Endogenous tagging of *pfvp1* with 3HA (hemagglutinin)

To modify the endogenous locus of *pfvp1* (PF3D7_1456800) via CRISPR/Cas9^23,24^, we made a template plasmid and two gRNA plasmids. Briefly, the *pfvp1* locus was tagged with 3HA and the regulatory elements (TetR-DOZI-aptamers)^25,26^ required to regulate the expression of *pfvp1*-3HA under the control of the small molecule aTc (anhydrotetracycline). For constructing the pMG75 template plasmid, we amplified two homologous regions (5HR and 3UTR) from WT genomic DNA using primers P1-P4. These two PCR products were cleaned and annealed together by an extension PCR method^54^. The assembled 3UTR+5HR fragment was digested with BssHII and BstEII, cloned into pMG75-3HA^55^, and sequenced using vector primers (P5-P6). These procedures resulted in the template plasmid, pMG75-PfVP1-3HA. For parasite transfection, the template plasmid was linearized with EcoRV and mixed with two circular gRNA plasmids.

For constructing gRNA plasmids, we selected the best two gRNA sequences using the Eukaryotic Pathogen CRISPR guide RNA Design Tool (http://grna.ctegd.uga.edu/) and cloned them individually into our NF-Cas9-yDHOH(-) plasmid^55^, bearing Cas9 from *Streptococcus pyogenes*. Cloning of gRNAs was carried out via NEB HiFiDNA Assembly as previously described^55^. Oligo sequences are listed as P7-P10. The cloned gRNAs were sequenced using a vector primer (P11).

#### 2) Endogenous tagging of *pfvp1* with mNeonGreen

We amplified the mNeonGreen sequence from pM2GT-Hsp101-mNeonGreen plasmid (kindly provided by Dr. Joshua Beck, Iowa State University) using P12-P13. The PCR product was digested and cloned into pMG75-PfVP1-3HA using SalI and ApaI sites. The replacement of 3HA with mNeonGreen was confirmed by sequencing using primers P12-P13.

#### 3) Complementing PfVP1 knockdown parasites with WT or mutant PfVP1 alleles

In the D10-PfVP1-3HA^apt^ line, we performed the second transfection to add back either a WT or mutated PfVP1 allele under a strong *P. falciparum* promoter (Cam, PF3D7_1434200). To this end, we first replaced the PfmtRL2 promoter in the pLN-RL2-hDHFR-3Myc construct^42^ with the Cam promoter^56^, yielding pLN-Cam-hDHFR-3Myc. We then amplified WT PfVP1 from genomic DNA using P16-17 and cloned it into this plasmid via AvrII and BsiWI sites, yielding pLN-Cam-hDHFR-PfVP1-3Myc. To make individual mutant PfVP1 alleles (D236A, D247A, D461A, L697A), we introduced each mutation into primers (P18-25), amplified two homologous fragments bearing the mutation, and assembled them into a full-length fragment using NEB HiFiDNA Assembly. Each of the assembled full-length mutant PfVP1 fragment was amplified, digested, and cloned into pLN-Cam-hDHFR-3Myc via AvrII and BsiWI, resulting in 4 mutant PfVP1 plasmids. All WT and mutant PfVP1 alleles were sequenced using the pLN vector primers (P26-P27).

#### 4) Cloning *Saccharomyces cerevisiae* inorganic pyrophosphatase (yIPP1) for complementation studies

The yeast IPP1 was amplified from *S. cerevisiae* genomic DNA using primers P28-29, cloned into pLN-Cam-hDHFR-3Myc via AvrII and BsiWI sites, and sequenced by primers (P26-P27).

#### 5) Cloning *Arabidopsis thaliana* VP1 (AVP1) for complementation studies

The AVP1 gene was amplified from p426CUP1-TcGFPAVP1^37^ using primers P30-31, cloned into pLN-Cam-hDHFR-3Myc via AvrII and BsiWI sites, and sequenced by primers (P26, P27, P32).

### 2, Plasmid construction for yeast studies (*Saccharomyces cerevisiae*)

#### 1) The yeast expression vectors were kindly provided by Dr. Kendal Hirschi from Baylor College of Medicine^37^

The plasmid p426CUP1-TcGFPAVP1 bears the copper inducible promoter CUP1, which drives the expression of the N-terminally tagged AVP1. The N-terminal tag, TcGFP, is the fusion of the first 28 aa of *Trypanosoma cruzi* VP1 (TcVP1) plus the full-length green fluorescent protein (GFP), which facilitates AVP1’s localization onto yeast vesicles^36^. The plasmid p426CUP1 contains nutritional markers including his3, trp1, leu2, and ura3.

#### 2) Cloning WT codon optimized PfVP1 for yeast expression

The scPfVP1 (synthetic codon optimized PfVP1 using *S. cerevisiae* codon usage tables) was synthesized by GeneWiz and cloned into a pUC plasmid (pUC-Kan-scPfVP1). The scPfVP1 sequence is also flanked by two homologous regions matching the ends of the p426CUP1-TcGFPAVP1 vector digested by BsrGI. The scPfVP1 sequence plus homologous regions was released from the pUC plasmid by SacII and cloned into p426CUP1-TcGFPAVP1 via NEB HiFiDNA Assembly. Replacement of AVP1 by scPfVP1 was verified by sequencing using primers P43-P45.

#### 3) Cloning mutant PfVP1 alleles for yeast expression

Using p426CUP1-TcGFPscPfVP1 as the template, we made 4 mutant PfVP1 alleles bearing mutations of D236A, D247A, D461A, and L697A, respectively. Using the similar mutagenesis strategy as described above, we introduced mutations into primers, amplified homologous fragments using primers (P33-P42), assembled the fragments into full-length sequences, and cloned them individually into p426CUP1, resulting in 4 mutant plasmids p426CUP1-TcGFPscPfVP1-D236A/D247A/D461A/L697A. All mutant PfVP1 alleles were sequenced using primers (P43-P45).

All primers used in this study were synthesized by GeneWiz and listed in **Supplementary file 1**. All cloned fragments were sequenced by Sanger sequencing (GeneWiz). All restriction enzymes were ordered from New England Biolabs.

### 3, Parasite culture, transfection, and knockdown studies

The 3D7-PfVP2KO (PfVP2 knockout) line was generated previously^21^. We used RPMI-1640 media supplemented with Albumax I (0.5%) to culture *P. falciparum* parasites in human O^+^ RBCs as previously described^42,55^. We transfected *P. falciparum* ring stage parasites (∼ 5% parasitemia) either with linearized or circular plasmid (∼ 50 µg) using a BioRad electroporator. Post electroporation, parasite cultures were maintained in proper drug selections, e.g., blasticidin (2.5 µg/mL, InvivoGen), WR99210 (5 nM, a kind gift from Jacobs Pharmaceutical), and anhydrotetracycline (aTc) (250 nM, Fisher Scientific). Parasite synchronization was performed with several rounds of alanine/HEPES (0.5M/10 mM) treatment. For knockdown studies, synchronized parasites were washed thrice with 1xPBS to remove aTc and diluted in fresh blood (1:10) to receive aTc (+) or (-) media.

### 4, Yeast culture, yeast lines and transformation

The *S. cerevisiae* strain BJ5459 was kindly supplied by Dr. Katrina Cooper from Rowan University^35^, which was originally created by^34^. This strain (*MAT*a, *his3*Δ*200, can1, ura3–52, leu2*Δ*1, lys2–801, trp1-289, pep4*Δ*::HIS3, prb1*Δ*1*.*6R*) lacks yeast vacuolar proteases PrA (proteinase A) and PrB (proteinase B). Yeast cultures were maintained at 30ºC either in YPD or Uracil drop-out medium. YPD medium contains 1% yeast extract (BP1422-500, Fisher Scientific), 2% peptone (20-260, Genesee Scientific), and 4% dextrose. Ura drop-out medium contains uracil minus complete supplement mixture (1004-010, Sunrise Science) and dropout base powder (1650-250, Sunrise Science). The latter has yeast nitrogen base (1.7 g/L), ammonium sulfate (5 g/L) and dextrose (20 g/L). Ura drop-out solid medium contains extra 2% agar. Yeast transformation was carried out using the Frozen-EZ Yeast Transformation II Kit (T2001, Zymo Research), according to manufacturer’s protocols.

### 5, Immunofluorescence analysis (IFA) and immuno-electron microscopy (Immuno-EM)

IFA was carried out as previously described^42,55^. Immuno-EM was performed at the Molecular Microbiology Imaging Facility at Washington University in St. Louis, MO. We used the following primary antibodies and dilutions: the HA probe (mouse, sc-7392, Santa Cruz Biotechnology; 1:300), the Myc probe (rabbit, 2278S, Cell signaling; 1:300), PfExp2 (rabbit, a kind gift from Dr. James Burns, Drexel University; 1:500), and PfPlasmepsin II (rabbit, Bei Resources, NIAID/NIH; 1:1000). We used fluorescently labeled secondary antibodies from Life Technologies (ThermoFisher Scientific) (anti-mouse or anti-rabbit, 1:300) or goat anti-mouse 18 nm colloidal gold-conjugated secondary antibody (Jackson ImmunoResearch Laboratories), as described previously^55^. Other details can be found^55^.

### 6, Western blot

Infected RBCs were lysed with 0.05% Saponin/PBS supplemented with 1x protease inhibitor cocktail (Apexbio Technology LLC) and protein was extracted with 2%SDS/62 mM Tris-HCl (pH 6.8) as previously described^55^. After protein transfer, the blot was stained with 0.1% Ponceau S/5% acetic acid for 5 min, de-stained by several PBS washes, and blocked with 5% non-fat milk/PBS. We used the following primary antibody dilutions: the HA probe (1:10,000), the Myc probe (1:8,000), and PfExp2 (1:10,000). We used HRP conjugated goat anti-mouse secondary antibody (A16078, ThermoFisher Scientific) at 1:10,000 or goat anti-rabbit HRP-conjugated secondary antibody (31460, ThermoFisher Scientific) at 1:10,000. Other steps followed the standard Bio-Rad Western protocols. For all Western samples, protein concentration was determined using the detergent tolerant Pierce™ BCA Protein Assay Kit (23227, ThermoFisher) according to the manufacturer’s protocols. Blots were incubated with Pierce™ ECL substrates and developed by the ChemiDoc Imaging Systems (Bio-Rad).

### 7, pH measurement using BCECF-AM (2’,7’-Bis-(2-Carboxyethyl)-5-(and-6)-Carboxyfluorescein, Acetoxymethyl Ester)

We measured the pH of saponin permeabilized parasitized RBCs using the pH-sensitive fluorescent dye (BCECF-AM) according to published protocols^15^. For each measurement, 0.5-1×10^^7^ parasitized RBCs were incubated with 4 µM BCECF-AM (B1170, ThermoFisher) and 0.02% Pluronic F-127 (p6867, ThermoFisher) in 16% hematocrit for 30 min at 37ºC. Pluronic F-127 was used to facilitate the diffusion of BCECF-AM across cellular membranes. In both aTc (+) and aTc (-) conditions, one aliquot of parasitized RBCs was incubated with Pluronic F-127 alone to serve as a negative control. After incubation, infected RBCs were permeabilized with 0.05%Saponin/PBS and washed twice with warm saline/glucose buffer (NaCl 125 mM, KCl 5 mM, MgCl2 1 mM, glucose 20 mM, HEPES 25 mM, pH 7.4). The pellet was resuspended in 1 mL warm saline/glucose buffer, transferred to a cuvette, and placed in the temperature-controlled chamber of a spectrofluorometer (Hitachi F-7000). The cell suspension was successively excited at 440 and 490 nm over 150 seconds and emitted fluorescence was measured at 535 nm. The ratio of fluorescence intensity excited by two wavelengths (490/440 nm) is a quantitative indicator of cellular pH.

To convert fluorescence intensity ratios to actual pH values, we calibrated pH measurement using the proton ionophore Nigericin according to published protocols^15^. Three aliquots of parasitized RBCs were incubated with BCECF-AM and Pluronic F-127 as described above, saponin treated, washed, and resuspended in a high K^+^ saline buffer (KCl 130 mM, MgCl2 1 mM, glucose 20 mM, HEPES 25 mM) at a pH of 6.8, 7.1 and 7.8, respectively. Nigericin (20 µM, AAJ61349MA, Fisher Scientific) was added to each aliquot of cell suspension before the sample was placed in the spectrofluorometer. Emitted fluorescence was recorded at 535 nm by dual-wavelength excitation at 440/490 nm as described above. Linear regression of fluorescence intensity ratios and pH values yielded an equation (regression coefficiency >0.99), which was used to calculate pH values of individual samples.

### 8, PPi extraction and measurement

PPi extraction was carried out following the published protocol with some modifications^57^. We used saline/glucose buffer (NaCl 125 mM, KCl 5 mM, MgCl2 1 mM, glucose 20 mM, HEPES 25 mM, pH 7.4) for saponin lysis and washes. At each timepoint from aTc (+) or (-) conditions, 2×10^^8^ parasitized RBCs (or uninfected RBCs as a control) were saponin lysed and washed 3 times to remove hemoglobin. The pellet was resuspended in 2 volumes of saline/glucose buffer, heated at 90ºC for 10 min to inactivate soluble pyrophosphatases, and saved at - 80ºC. The samples were thawed and undergone 4 cycles of freezing/thawing between dry ice (10 min) and 37ºC (∼ 2 min). They were sonicated for 40 min at 4ºC in a water bath sonicator (Fisher). After sonication, samples were spun down at 13,000 rpm for 10 min. The supernatants were saved for PPi measurement. The pellets were solubilized with 2%SDS/62 mM Tris-HCl (pH 6.8) overnight for protein quantification.

PPi was measured with a PPi fluorogenic sensor from Abcam (ab179836) (the chemical identity of this sensor was not released by the manufacturer). Briefly, 2 µL of each supernatant as extracted above was added into a 50 µL assay buffer containing 1:1000 diluted PPi fluorogenic sensor in a black plate. The mixture was incubated in the dark for 20-30 min and read by Tecan infinite 200 pro at 470 nm with excitation at 370 nm. A PPi standard curve was generated to determine PPi concentrations in samples.

### 9, SYBR Green Assays

We performed SYBR green assays as previously published^55^. Compounds used in this study included Bafilomycin A1 (NC1351384, Cayman Chemical), chloroquine (AC455240250, Fisher Scientific), atovaquone (A7986, MilliporeSigma), and artemisinin (a kind gift from Dr. Jianping Song at Guangzhou University of Chinese Medicine, China). In brief, drugs were serially diluted (3-fold) in 96 well plates in regular medium. Parasites from aTc plus culture (0.5% ring at 4% hematocrit) were washed several times with PBS and resuspended in various concentrations of aTc (20, 10, 2, or 0 nM) and incubated with diluted drugs, yielding final concentrations of aTc at 10, 5, 1, and 0 nM. Data was analyzed by GraphPad Prism6.

### 10, Yeast vesicle isolation

We followed published protocols to purify yeast vesicles expressing various VP1 proteins^37^. In brief, from the transformed plate, one colony was picked and inoculated into 5 mL of drop-out medium on Day 1. Day 2, the culture was diluted in 1:33 and grown overnight in 100 mL of drop-out medium. Day 3, the culture was diluted in 1 L of YPD (starting OD_600_=0.05) and grown to reach OD_600_ 0.8 (typically, < 12 h). The culture was then induced with 3 mM of CuSO_4_ for 3-4 h to reach OD_600_ between 1-1.3.

After induction, the yeast cells were pelleted, washed with deionized water and spheroplast buffer (1.2 M Sorbitol, 100 mM KH_2_PO_4_, pH 7.0), and weighed. The pellet was then resuspended in five volumes of spheroplast buffer containing 10 mM DTT and 1% glucose (both freshly added). In this mixture, per gram of wet yeast, 5 mL of Zymolyase 20T (120494-1, AMSBIO) at 5 mg/ml in 10 mM Na_2_HPO_4_, 50% Glycerol was added. This mixture was incubated at 30°C with rotating for 2 h to digest the yeast cell wall. After digestion, the yeast pellet was washed twice with spheroplast buffer, resuspended in Lysis Buffer A (10 mM MES-Tris, 0.1 mM MgCl_2_, pH 6.9, 12% Ficoll 400 (AAB2209518, Fisher Scientific, added fresh)). The mixture was dounced 15-25 times on ice and pelleted. The supernatant was transferred to an ultracentrifuge tube and overlayed with layers of Lysis Buffer A with 12% Ficoll and Buffer B with 8% Ficoll (AAB2209518, Fisher) and centrifuged at 28,500 rpm for 45 min (Beckman 42.1). Afterwards, the wafer clump was collected using a pipette tip and resuspended in 3 mL of 2x Buffer C and mixed with 3 mL of 1x Buffer C (20 mM MES-Tris, 10 mM MgCl_2_, 50 mM KCl, pH 6.9) and pelleted in the ultracentrifuge (Beckman 50Ti). The resulting pellet was then resuspended in 200 µL of 1x Buffer C with 10% glycerol, aliquoted into microcentrifuge tubes and flash frozen in an ethanol dry ice bath before storage at -80°C.

### 11, ACMA pH Quenching Assay

We measured proton pumping activities of VP1 in isolated yeast vesicles using the ACMA Fluorescence Quenching Assay^37^. ACMA stands for 9-amino-6-chloro-2-methoxyacridine (A1324, ThermoFisher Scientific). For each measurement in the spectrofluorometer (Hitachi F-7000), 30 µg of vesicles were added to the 1 mL of transport buffer (100 mM KCl, 50 mM NaCl, and 20 mM HEPES) in the presence of 1 µM of ACMA, 3 mM MgSO_4_, 1 mM of Na_2_PPi and 1 µM of Bafilomycin A1 (inhibitor of the yeast V-type ATPase). The reaction was monitored for 15 minutes to observe any decrease in fluorescence (excitation 410 nm, emission 490 nm). Afterwards, 10 µM of Nigericin is added to the solution and monitored for 3 min to see if the fluorescence could be restored.

### 12, Pyrophosphatase activity measurement

The release of Pi by pyrophosphatase activity of VP1 proteins in isolated yeast vesicles was measured using the P_i_Per ™ Phosphatase Assay Kit (P22061, ThermoFisher Scientific), according to manufacturer’s protocols. The Pi derived from PPi hydrolysis is coupled to three enzymatic reactions to convert a nonfluorescent compound (amplex red) to fluorescent resorufin. Fluorescence was detected by Tecan infinite 200 pro at 565 nm with excitation at 530 nm. A Pi standard curve was generated to determine Pi concentrations in the samples.

## Acknowledgements

We thank members of the Ke lab, Swati Dass, Neeta Shadija, and Dr. Maruthi Mulaka, for technical assistance. We thank members of the Dr. Akhil Vaidya’s lab at Drexel University for constructive discussions and Dr. Michael Mather, Ian Lamb, and Swaksha Rachuri for editing the manuscript. We thank Dr. Kendal Hirschi’s lab (Baylor College of Medicine) and Dr. Katrina Cooper’s lab (Rowan University) for providing yeast plasmids and strains. We thank Dr. James Burn (Drexel University), Dr. Daniel Goldberg (Washington University St Louis), and Dr. Joshua Beck (Iowa State University) for providing antibodies and plasmids. We thank Dr. Jacquin Niles (Massachusetts Institute of Technology) and Dr. Sean Prigge (Johns Hopkins University) for providing the knockdown tools. We thank Dr. Wandy Beatty (Washington University St Louis) for performing immune-EM studies. This work was supported by a Career Transition Award from NIH/NIAID (K22AI127702) and a R21 grant from NIH/NIAID (1R21AI156735) to Dr. Hangjun Ke.

## Author contributions

O.S., L.L., and H.K. performed all experiments. T.M.F generated the modeled PfVP1 structures. H.K. and J.Z. designed all experiments. H.K. and O.S. wrote the manuscript, which was reviewed and edited by all other authors.

## Data availability

All data generated in this study have been included in the main and supplementary figures and in the source data files.

## Additional files

Supplementary files

Supplementary file 1. Primers used in this study.

Transparent reporting form

## Figure Legend

**Figure supplement 1. RNA-seq data of pyrophosphatases and proton pumps in *P. falciparum***.

Transcription data was retrieved from PlasmoDB (deposited by Bartfai *et al*.^16^) and plotted. PfVP1, *Plasmodium falciparum* vacuolar pyrophosphatase 1; PfVP2, *Plasmodium falciparum* vacuolar pyrophosphatase 2; sPPase, *Plasmodium falciparum* soluble pyrophosphatase; V-type ATPase subunit B and subunit A. FPKM stands for fragments per kilobase of transcript per million mapped reads.

**Figure supplement 2. Genetic tagging of PfVP1 via CRISPR/Cas9**.

A, Model of CRISPR/Cas9 mediated gene editing. The genetic locus of PfVP1 was tagged at the C-terminal with epitopes and aptamer repeats. Blasticidin deaminase served as the transfection marker. HR, homologous region. UTR, untranslated region. B, Model of aTc regulated conditional knockdown. Without aTc, the negative regulator, TetR-DOZI fusion protein, binds to the secondary structure of aptamers and protein translation is turned off. With aTc, the transcript is freed from TetR-DOZI and translated. aTc, anhydrotetracycline. TetR-DOZI, tetracycline repressor and development of zygote inhibited. C, Genotyping of Pf3D7VP2KO-VP1-3HA^apt^ by PCR. 5’int, 5’ integration. 3’int, 3’ integration. Star, a non-specific band. Primer positions are shown in A.

**Figure supplement 3. Localization of PfVP1 via Immunoelectron microscopy**.

Representative immuno-EM images of Pf3D7VP2KO-VP1-3HA labeled with anti-HA and gold-conjugated secondary antibodies. Black arrows indicate PVM, parasitophorous vacuolar membrane, and PPM, parasite plasma membrane. Red arrows indicate nuclear or cytosolic signals of PfVP1. Fv, food vacuole. Scale bars, 200 nm.

**Figure supplement 4. Co-localization of PfVP1 and the food vacuole via Immunofluorescence assay**. DAPI stains nuclei. PfVP1 was detected by anti-HA and Alexa Fluor™ 488 anti-mouse secondary antibodies. The food vacuole was detected by anti-PfPlasmepsin II^28^ and Alexa Fluor™ 568 anti-rabbit secondary antibodies. T, trophozoite. S, schizont. Representative images of n=50 parasites of each stage are shown here.

**Figure supplement 5. Localization of VP1 proteins in *S. cerevisiae***.

VP1 is N-terminally tagged with the localization peptide of TcVP1 (*Trypanosoma cruzi*) and GFP. Live microscopy showed localization of PfVP1 or AVP1 on the yeast’s vacuole. The yeast cell transformed with a negative control plasmid showed a fuzzy background. Representative images of n=30 yeast cells in each condition are shown here.

**Figure supplement 6. PfVP1 is highly expressed and essential for the ring stage and the transition to trophozoite stage**.

A, Western blot of Pf3D7VP2KO-VP1-3HA parasites after aTc removal for 24 and 48 h. PfVP1 was detected by anti-HA and anti-mouse HRP conjugated secondary antibodies. PfExp2 served as a loading control. The Ponceau stained membranes were shown at the bottom. Note the difference of loading in two blots, ∼ 1µg (left) versus ∼ 10 µg (right). **, when a small amount of total protein lysate was loaded, only aggregated forms of PfVP1 with high molecular weights were detected by Western blot. B, Parasitemia of Pf3D7VP2KO-VP1-3HA after aTc removal for 24, 48, 72 and 96 h. *, the arrested ring stage parasites were not counted at 96 h. This experiment was repeated three times.

**Figure supplement 7. Arrested ring stage parasites can poorly progress when aTc is restored**

A, Giemsa-stained images show parasite morphologies after aTc was added back for 24 and 48 h after it was previously removed for 96 h. Scale bars, 5 µm. B, Parasitemia of the addback culture for 24 and 48 h. This experiment was repeated four times.

**Figure supplement 8. Localization and expression of PfVP1 mutant alleles**.

A, Localization of PfVP1 mutant alleles in *P. falciparum*. The mutant PfVP1 protein was detected by anti-Myc and Alexa Fluor™ 488 anti-rabbit antibodies. Scale bar, 5 µm. Representative images of n=30 parasites of each condition are shown here. B, Expression of PfVP1 proteins by Western blot. From ∼3 µg of total protein lysate from each parasite line, the endogenous PfVP1 was detected by anti-HA and anti-mouse HRP secondary antibodies. **, when a small amount of total protein lysate was loaded, only aggregated forms of PfVP1 with high molecular weights were detected by Western blot. The same blot was re-probed with anti-PfExp2 antibody to show loading controls. In another blot, from ∼30 µg of total protein lysate from each parasite line, the episomal mutant PfVP1 protein was detected by anti-Myc and anti-rabbit HRP secondary antibodies. These experiments were repeated two times. C, Quantification of parasite morphologies of different mutant PfVP1 lines grown in aTc (-) media for 96 h. The percentage of ring or trophozoite stage was determined by counting 1000 infected RBCs in each condition. This experiment was repeated three times. D, Localization of codon optimized PfVP1 mutant alleles in *S. cerevisiae* via live microscopy. Each mutant PfVP1 protein was N-terminally tagged with the localization peptide of TcVP1 (*Trypanosoma cruzi*) and GFP. Representative images of n=30 yeast cells of each condition are shown here.

