## Supplementary Information for "Malaria parasites utilize pyrophosphate to fuel an essential proton pump in the ring stage and the transition to trophozoite stage" (from another version of this preprint: version 2, 2021-11-15)

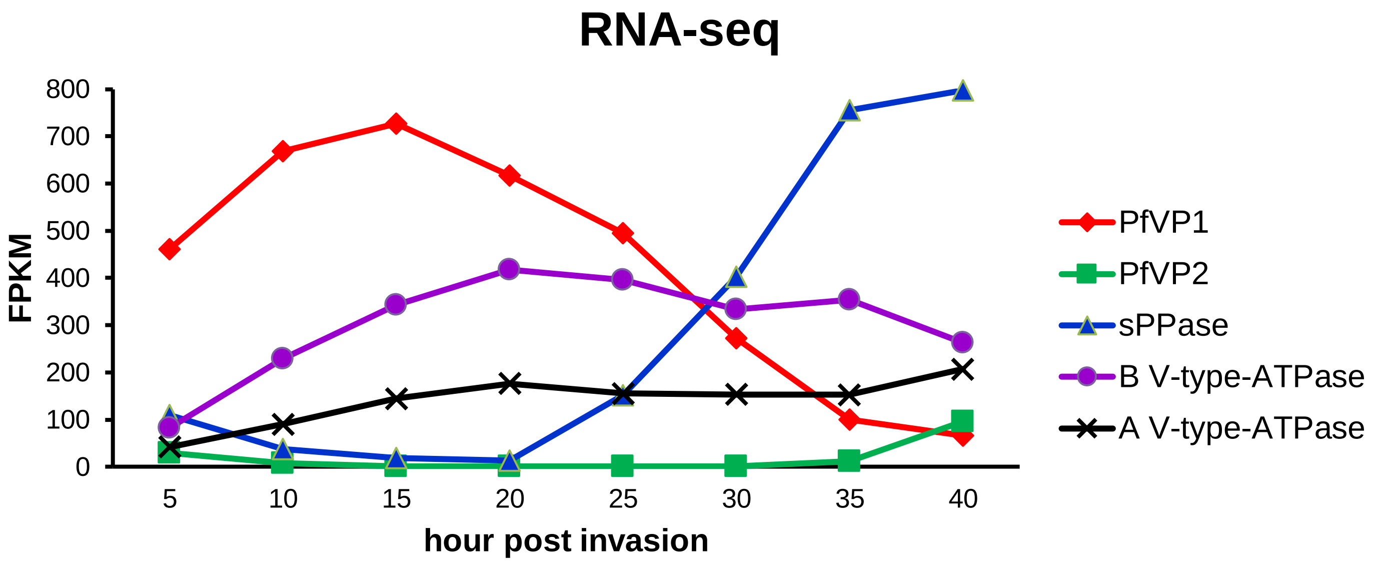


**Fig S1.** RNA-seq data of pyrophosphatases and proton pumps in *P. falciparum*.

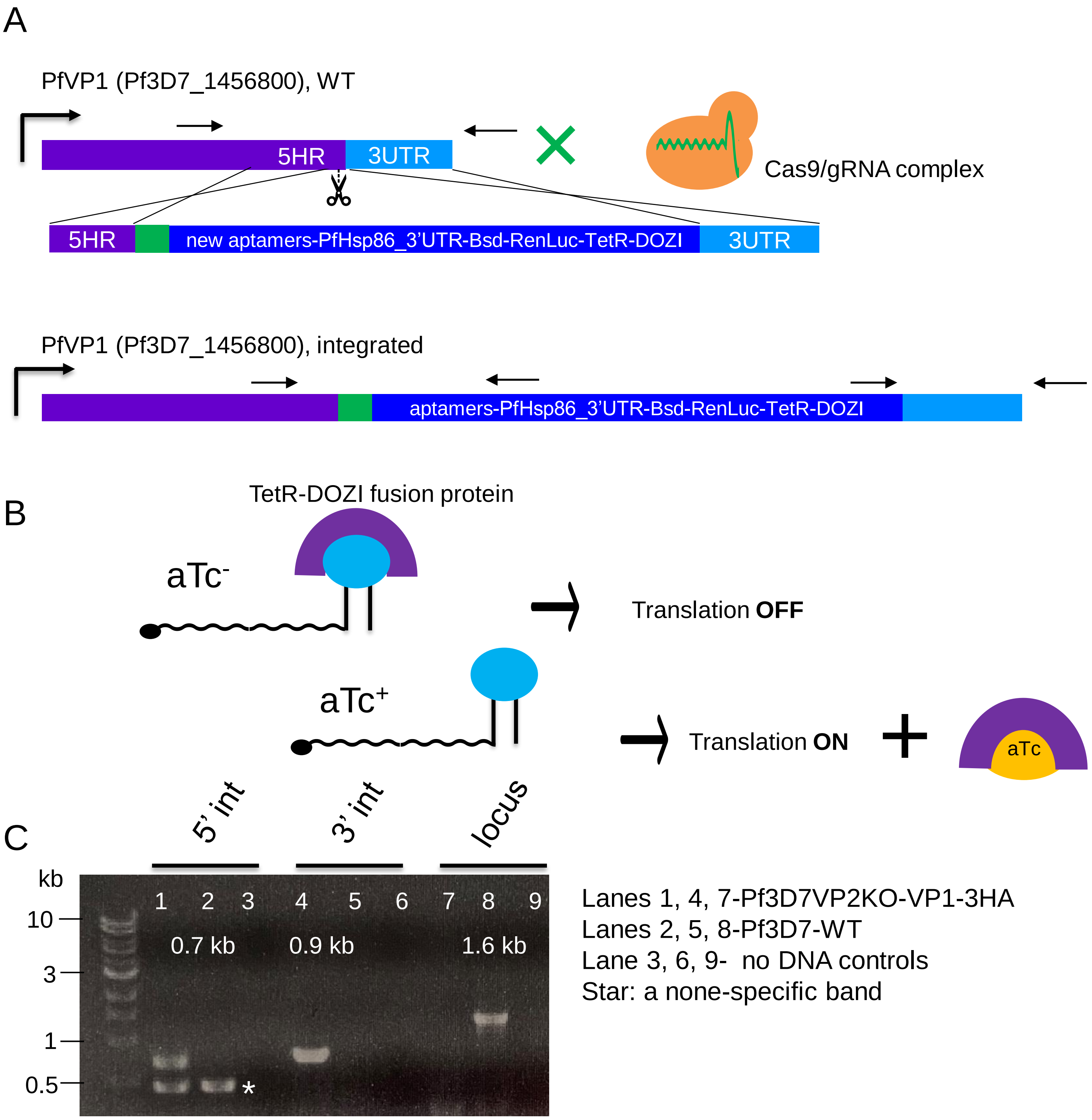


**Fig S2.** Genetic tagging PfVP1 via CRISPR/Cas9

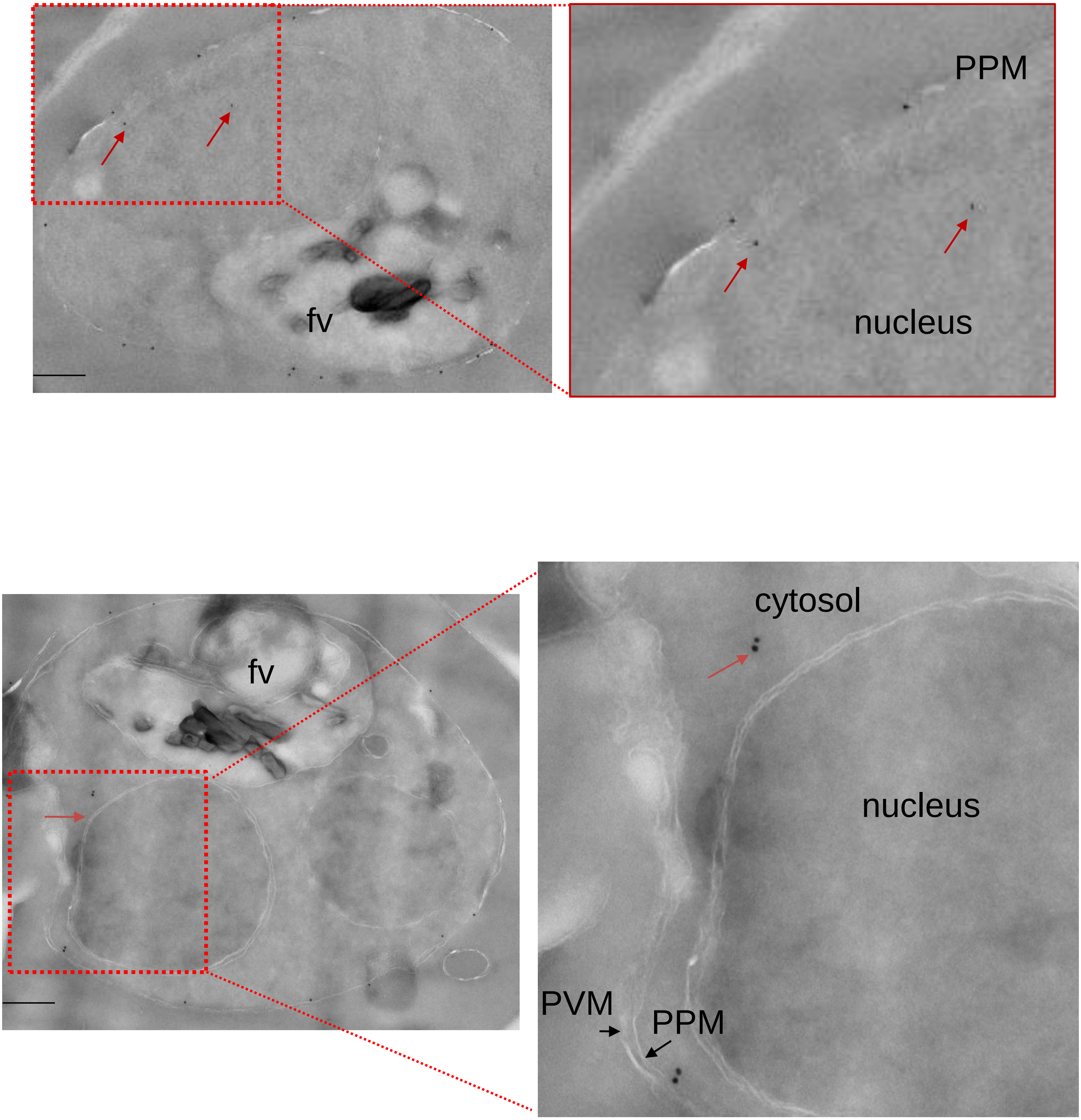


**Fig S3.** Localization of PfVP1 by Immunoelectron microscopy

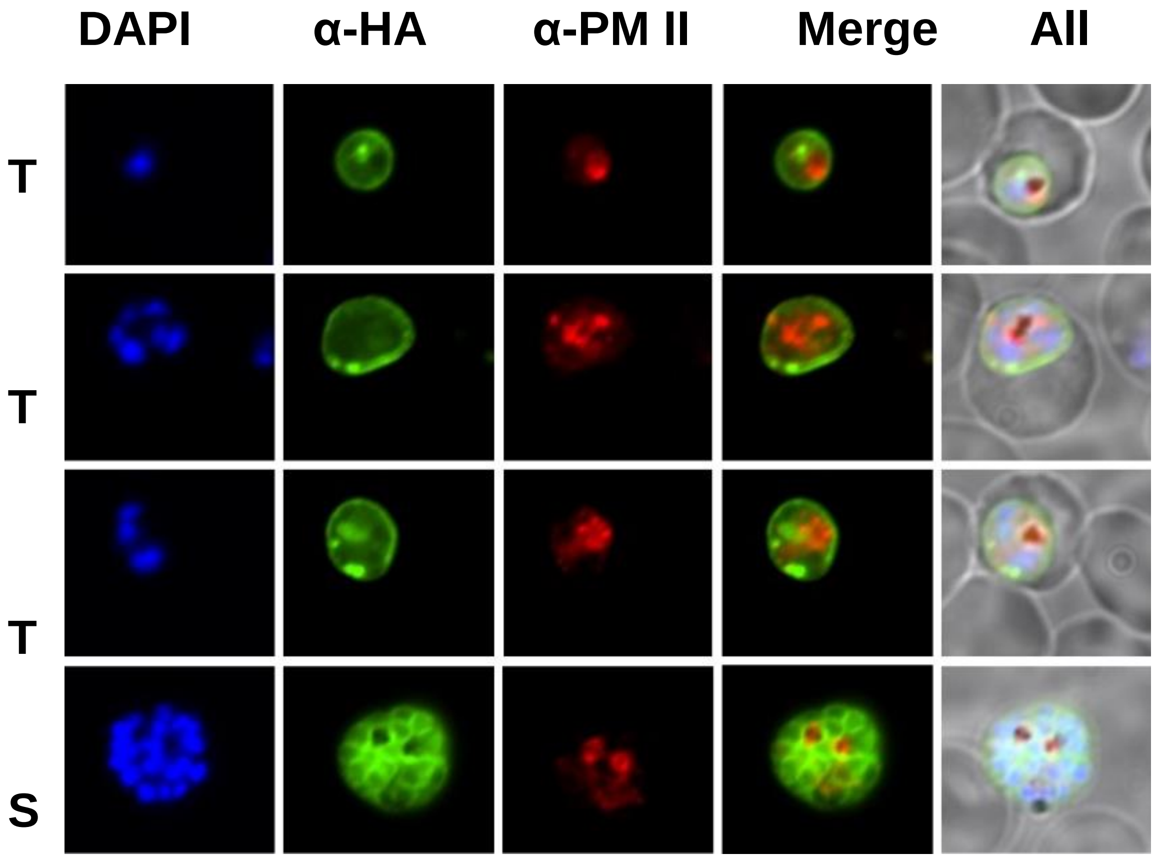


**Fig S4.** Co-localization of PfVP1 and Plasmepsin II via Immunofluorescence assay

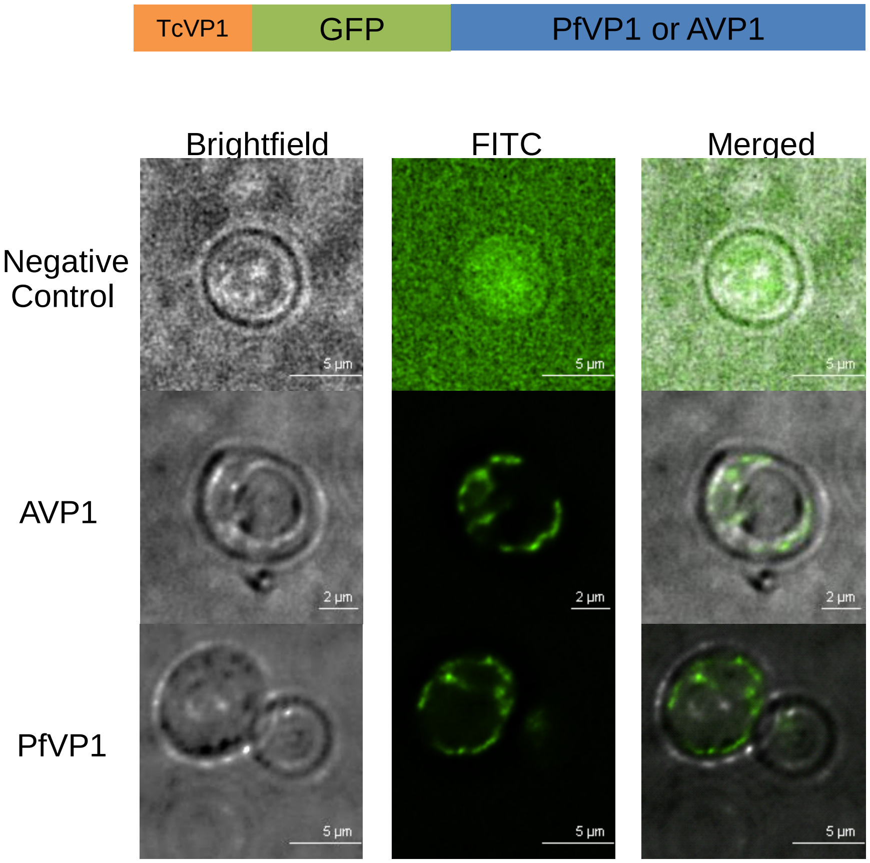


**Fig S5.** Localization of VP1 proteins in *S. cerevisiae*

A

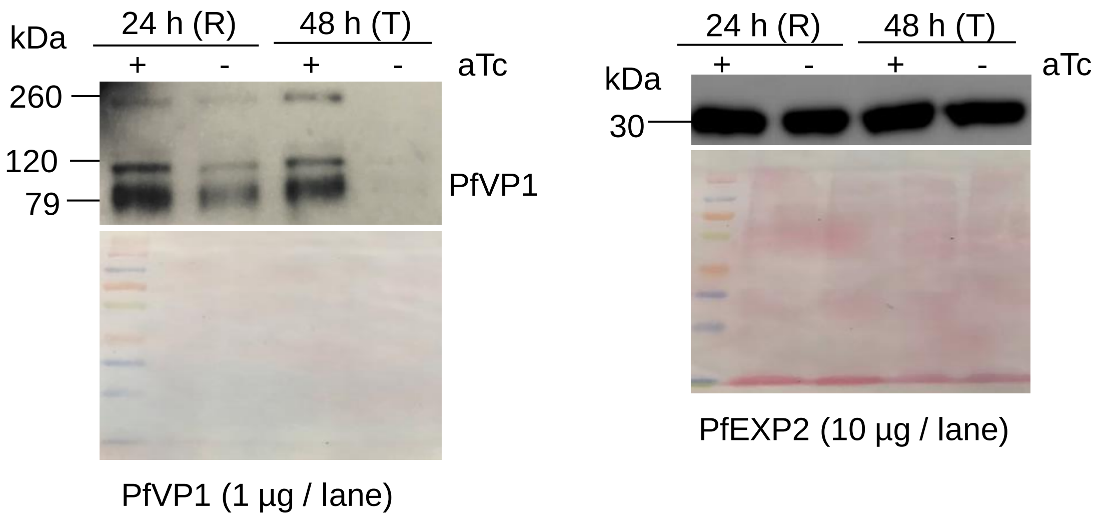


B

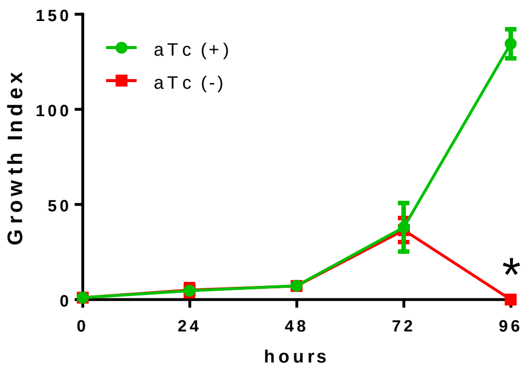


**Fig S6.** PfVP1 is highly expressed and essential for ring to trophozoite transition

A

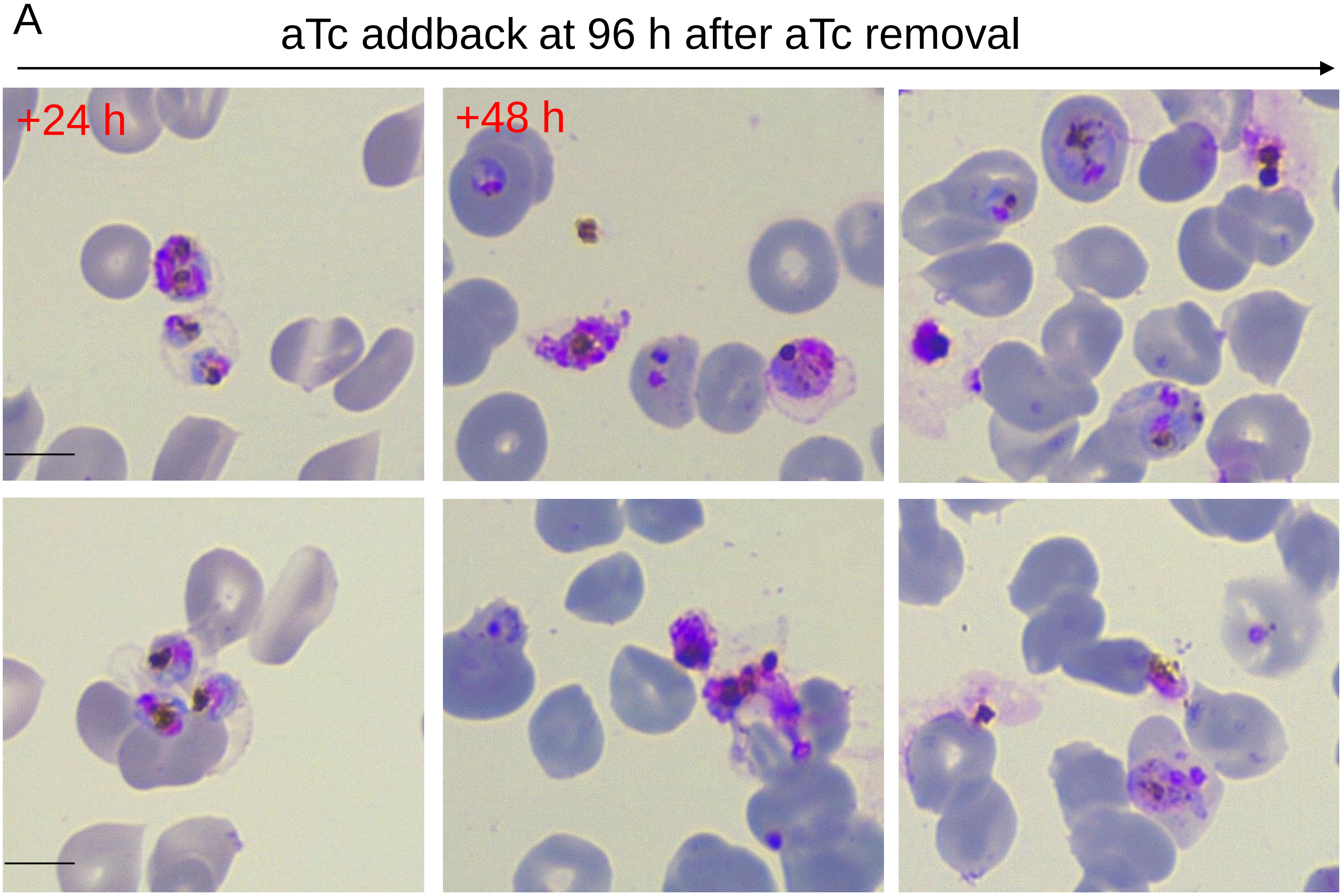


B

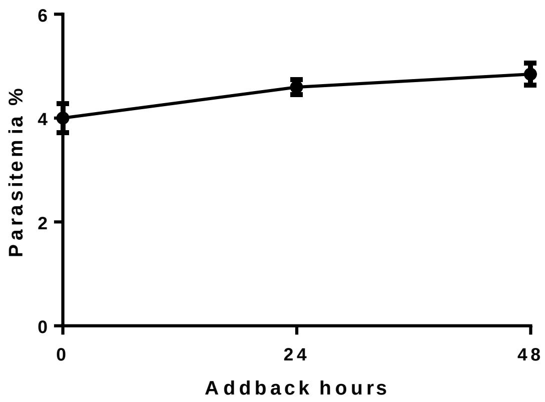


**Fig S7.** Arrested ring stage parasites can poorly progress when aTc is restored

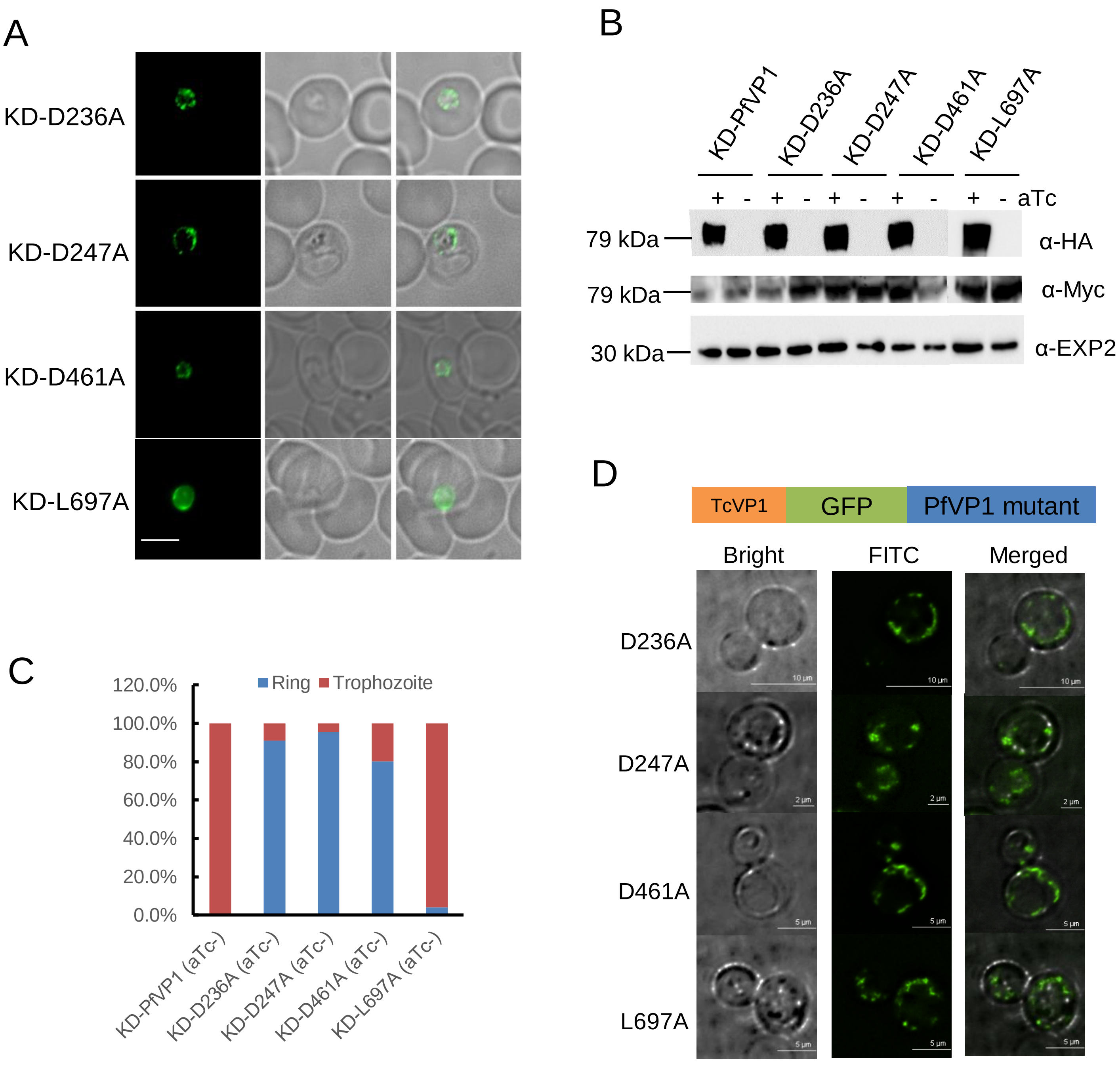


**Fig S8.** Localization and expression of PfVP1 mutant proteins
